# Brain activity fluctuations propagate as waves traversing the cortical hierarchy

**DOI:** 10.1101/2020.08.18.256610

**Authors:** Yameng Gu, Lucas E. Sainburg, Sizhe Kuang, Feng Han, Jack W. Williams, Yikang Liu, Nanyin Zhang, Xiang Zhang, David A. Leopold, Xiao Liu

**Affiliations:** Department of Biomedical Engineering, The Pennsylvania State University, University Park, PA, 16802, USA; The Huck Institutes of the Life Sciences, The Pennsylvania State University, University Park, PA, 16802, USA; College of Information Sciences and Technology, The Pennsylvania State University, University Park, PA, 16802, USA; Neurophysiology Imaging Facility, National Institute of Mental Health, National Institute of Neurological Disorders and Stroke, and National Eye Institute, National Institutes of Health, Bethesda, MD, 20892, USA; Section on Cognitive Neurophysiology and Imaging, Laboratory of Neuropsychology, National Institute of Mental Health, National Institutes of Health, Bethesda, MD, 20892, USA; Institute for Computational and Data Sciences, The Pennsylvania State University, University Park, PA, 16802, USA

**Keywords:** infra-slow propagating activity, cortical hierarchy, multimodal neuroimaging, global signal

## Abstract

The brain exhibits highly organized patterns of spontaneous activity as measured by resting-state fMRI fluctuations that are being widely used to assess the brain’s functional connectivity. Some evidence suggests that spatiotemporally coherent waves are a core feature of spontaneous activity that shapes functional connectivity, though this has been difficult to establish using fMRI given the temporal constraints of the hemodynamic signal. Here we investigated the structure of spontaneous waves in human fMRI and monkey electrocorticography. In both species, we found clear, repeatable, and directionally constrained activity waves coursed along a spatial axis approximately representing cortical hierarchical organization. These cortical propagations were closely associated with activity changes in distinct subcortical structures, particularly those related to arousal regulation, and modulated across different states of vigilance. The findings demonstrate a neural origin of spatiotemporal fMRI wave propagation at rest and link it to the principal gradient of resting-state fMRI connectivity.

## Introduction

The human brain represents about only 2% of the total body weight but accounts for ∼20% of the total energy budget, and a majority (∼95%) of brain energy is consumed by intrinsic brain activity at rest (1, 2). This budget allocation is consistent with a highly organized nature of fMRI signals collected in the resting state, which are being widely used for inferring functional brain connectivity in health and disease (3, 4). The study of resting-state fMRI (rsfMRI) dynamics has suggested that the highly structured rsfMRI connectivity, i.e., correlations, may arise from transient fMRI co-activations caused by event-like brain activity (5–8), which were also to show systematic transitioning patterns (9, 10). Consistent with these findings, the propagating structures have been found in rsfMRI by using a template-refining approach to extract repeated quasi-periodic patterns (11–13) or by decomposing rsfMRI lag structures to recover lag threads (14, 15). These propagating structures contribute significantly to rsfMRI connectivity (14, 16) and appear sensitive to brain state changes and diseases (17–19), and could be the key to understanding the functional role of intrinsic brain activity (20).

Nevertheless, to date the neural origin of the rsfMRI propagations remains elusive. The study of propagating activity using fMRI faces a serious issue due to the spatial heterogeneity of hemodynamic delays (21). A series of recent studies have shown that a systemic low-frequency oscillation of blood signals, which can be recorded at peripheral sites such as fingertips and toes, induces systematic rsfMRI delays across brain regions that are consistent with the blood transit time through the cerebrovascular tree (22–26), suggesting a potential contribution of hemodynamic delays to apparent rsfMRI propagations. On the other hand, simultaneous fMRI-electrophysiology recordings in rats have provided clear evidence for the co-modulation of neural activity with the rsfMRI propagations (12), which is however insufficient to prove the neural origin of the “propagation” per se. In other experimental setups, spatially propagating waves have been observed among neural populations, e.g., wide-field optical imaging of voltage-sensitive dye or calcium in mice. However, such waves are difficult to compare with those measured with fMRI, as they are usually local (millimeters) and on a rapid time scale (<1 s) (27, 28). Most recently, globally propagating waves on the seconds timescale (∼5 seconds) and the lag structure were observed in mice using calcium imaging and suggested to account for resting-state hemodynamic connectivity (18, 29). However, it remains unclear whether similar resting-state infra-slow propagations over the entire cortex are present in neural signals of awake primates, and if so, whether and how are they similar to the propagating activity in human rsfMRI. There is also a lack of a detailed characterization of the infra-slow propagating activity, including its trajectories and subcortical involvements. Furthermore, although the contribution of these slow propagations to rsfMRI connectivity has been demonstrated (14–16, 29, 30), it remains unclear whether they can be linked to any specific component of rsfMRI connectivity. Both the major quasi-periodic pattern (11) and the latency projection of the lag structure (14, 15) showed a distinct contrast between the default-mode network and sensory/motor regions. This contrast superficially resembles the so-called *principal gradient* of the brain’s spontaneous activity, which has been derived by embedding the rsfMRI connectivity matrix into a low-dimensional space (31, 32). This coincidence raises two important questions. First, is the direction of spatiotemporal fMRI propagation fundamentally linked to the reported rsfMRI connectivity gradient? And second, do the large-scale propagating waves reflect an underlying pattern of electrophysiological activity following the same trajectory?

To address these questions, this study combines human rsfMRI and monkey electrophysiology to study the infra-slow propagating brain activity. We developed a data-driven method to detect single propagating instances and map their propagating trajectories. The application of this method to high-resolution rsfMRI data of Human Connectome Project (HCP) revealed global propagations mostly in two opposite directions along a axis strikingly similar to the principal gradient of rsfMRI connectivity (31). The application of the same method to a large-scale electrocorticography (ECoG) recording from monkeys revealed very similar cross-hierarchy propagations between the lower- and higher-order brain regions, which are present most strongly at the gamma-band (42–95 Hz) power signals. Close inspection of the global rsfMRI propagations suggests local, embedded propagations within sensory modalities proceed in the opposite direction of the global propagation. More importantly, these cortical propagations are accompanied by sequential co-activation/de-activation in specific subcortical structures, particularly those related to arousal regulation. Consistent with this finding, the temporal dynamics of the infra-slow propagating activity are significantly modulated across brain states of distinct vigilance. Taken together, the study demonstrates a characteristic pattern of spontaneous, slowly propagating activity across the cortical hierarchy in humans and nonhuman primates, maps its detailed trajectories and associated subcortical changes, demonstrates its brain-state dependency, and also links it to the principal gradient of the rsfMRI connectivity.

## Results

### Infra-slow propagations along the principal gradient (PG) of rsfMRI connectivity

We first examined and characterized infra-slow propagating activity in rsfMRI signals using data from 460 HCP subjects. Simple visual inspection of pre-processed signals suggested clear propagating activity that often coursed from higher-order cognitive areas, mostly the default-mode network, to lower-order sensory/motor regions, a direction similar to the principal gradient (PG) of rsfMRI connectivity, or in an opposite direction. Following previous work (31), the principal gradient was obtained by decomposing rsfMRI connectivity data using a data-driven method, and it reveals a gradient of functional connections that aligns well with the cortical hierarchy defined both anatomically and functionally (31). To visualize the propagating activity in a 2-D representation, we projected the rsfMRI signals onto the principal gradient direction (**Fig. 1A**) to generate time-position graphs (**Fig. 1B**). It was immediately noticed that local rsfMRI peaks at various principal gradient positions tended to cluster together in time and form continuous bands, some of which were tilted and propagated either from the sensory/motor to the default-mode network or in an opposite direction (**Fig. 1B**, upper), which we will refer to as the bottom-up and top-down propagations henceforth. In contrast, projecting the same signal onto a few other directions, including one obtained by randomly rotating the principal gradient map on brain surface (**Fig. 1B**, bottom), revealed straightly vertical bands, suggesting an absence of propagating behavior along these directions. To locate and quantify single propagating instances, we cut the rsfMRI signals into time segments based on troughs of the global mean signal, which successfully separated the bands in the time-position graphs (**Fig. 1B**). For each segment, we correlated the relative timing of local rsfMRI peaks with their relative position along different directions. A high time-position correlation would suggest a propagation along the corresponding direction (**Fig. 1C** and **Animations 1-2**). We then computed and summarized the time-position correlations for all rsfMRI segments showing a global involvement (defined as the global signal peak exceeding a threshold established using a null model, see **Methods** and **Fig. S1** for the details), which account for 58.80 % of the total segments. The time-position correlations of local peaks for the principal gradient direction showed a non-Gaussian, bimodal (*p* = 0.012, Hartigan’s dip test) distribution with a relatively larger peak for positive correlations, indicating significant propagations along this axis, particularly in the bottom-up direction (**Fig. 1D**, the left column). In contrast, they showed Gaussian-like, single-mode distributions for the other four control directions (**Fig. 1D**, the right four columns), which are significantly different from the principal gradient distribution (*p* = 0 for all four control directions, two-sample Kolmogorov-Smirnov test). This is even true for the second gradient of rsfMRI connectivity (31) that showed a strong motor-to-visual contrast (**Fig. 1D**, the second column), suggesting a lack of propagations along this direction in these rsfMRI time segments. All these results suggest the existence of significant rsfMRI propagating activity along the principal gradient direction.

**Fig. 1.**
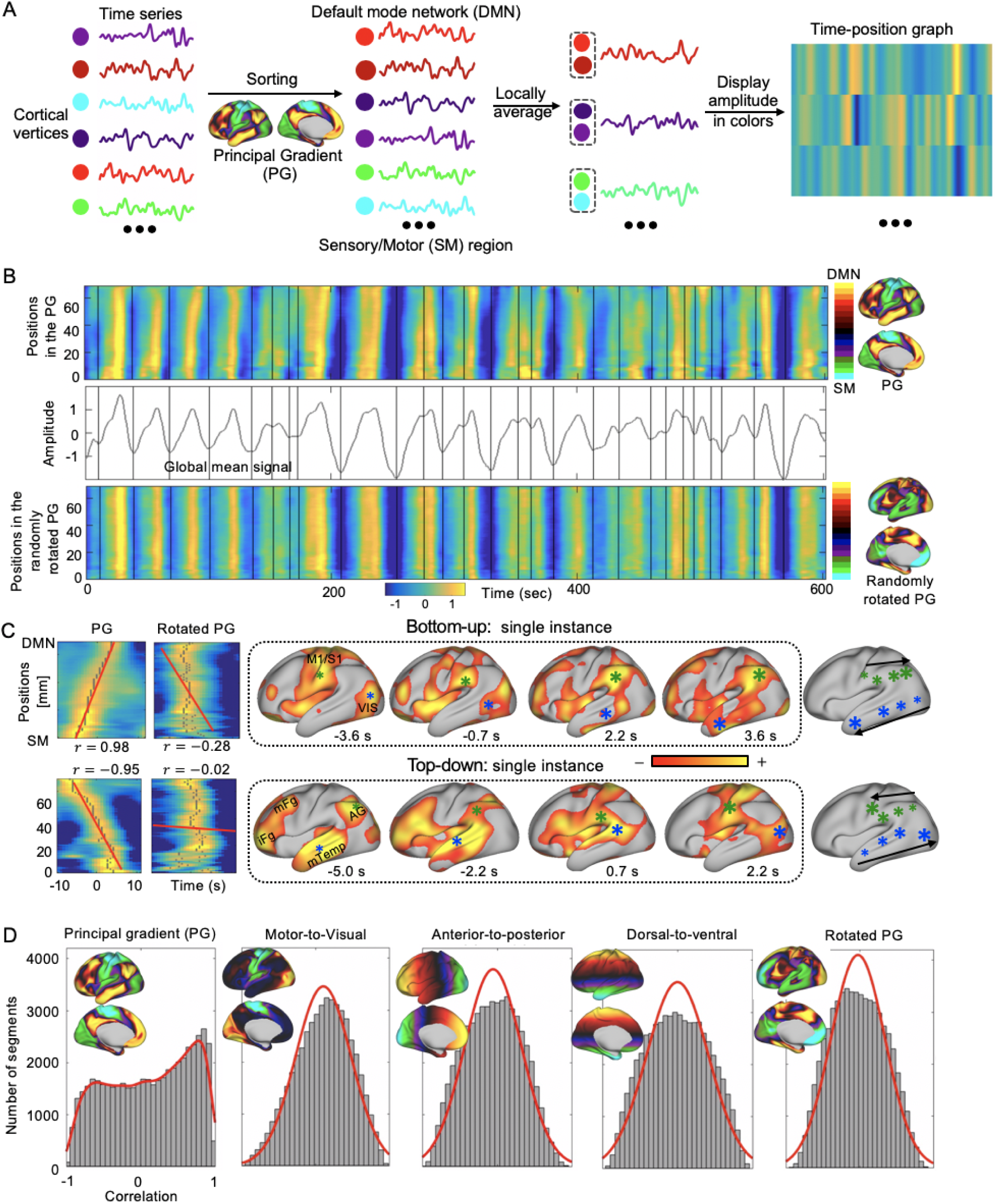
RsfMRI propagations along the principal gradient (PG) of rsfMRI connectivity. (A) Illustration of the steps for projecting rsfMRI signals onto a specific direction, e.g., the principal gradient, to produce a time-position graph. The principal gradient was computed in a previous study (31) by applying the diffusion mapping to the averaged connectome matrix and indicted a transition across brain hierarchies from the SM regions to the DMN. (B) The time-position graphs for the principal gradient and a control direction from a representative subject show clear bands, which are tilted only in the principal gradient time-position graph. The bands can be well separated by cutting rsfMRI segments according to troughs (black vertical lines) of the global mean signal. The vertical axis of the time-position graph represents the cortical distance from core regions of the DMN to the SM on the principal gradient (31). (C) Two exemplary rsfMRI segments with propagations in two opposite directions along the principal gradient, i.e., the DMN-to-SM (i.e., top-down) and SM-to-DMN (i.e., bottom-up) propagations. The relative timing of the local rsfMR peaks show a significant correlation with their positions along the principal gradient but not in the control direction (left), and the propagating activity can be viewed on brain surface (right). Gray dots in the time-position graphs indicate the local rsfMRI peaks and the red lines are the regression lines for their time-position relationship. The time is with respect to the global mean peak for the segment. Green and blue asterisks mark approximately spatial peaks with a larger size representing a later time point and the timing being denoted by the direction of the black arrows. (D) The distributions of the time-position correlations of rsfMRI segments in five different directions. The principal gradient direction is associated with a clear bimodal distribution that is significantly different from those of the motor-to-visual, the anterior-to-posterior, the dorsal-to-ventral, and the rotated principal gradient directions (*p* = 0 for all four control directions, two-sample Kolmogorov-Smirnov test). Abbreviation: PG, principal gradient; SM, sensory/motor; DMN, default mode network; VIS, visual cortex; M1, primary motor cortex; S1, primary somatosensory cortex; iFG, inferior frontal gyrus; mFG, middle frontal gyrus; AG, angular gyrus; mTemp, middle temporal cortex.

### The cortical hierarchical axis is the dominant direction of rsfMRI propagations

We then developed a data-driven method to obtain main trajectories of rsfMRI propagations without setting a priori direction. The existing quasi-periodic pattern method was designed to find repeated spatiotemporal patterns that are not necessarily propagations (12), whereas the lag structure/thread method relied on session-based quantification of temporal lags in which the delays caused by anti-direction propagations may cancel each other (14, 15). We show that the propagations can cause systematic delays of local peaks and thus a significant time-position relationship along the propagating direction (**Fig. 1C**). To identify the major propagating direction in a data-driven way, we derived a delay profile for each rsfMRI segment by computing the relative delay of local peaks with respect to the global mean peak, and then applied a singular value decomposition to extract the principal components of all the delay profiles, which are expected to represent main trajectories of propagating activity (**Fig. 2A**). The application of this method to synthesized data containing simulated propagating structures successfully recovered two propagating directions (**Fig. 2B**). In comparison, the diffusive embedding method, which was employed to derive the principal gradient of rsfMRI connectivity (31), recovered the more frequent propagation but not the other one (**Fig. 2** and **Fig. S2**).

**Fig. 2.**
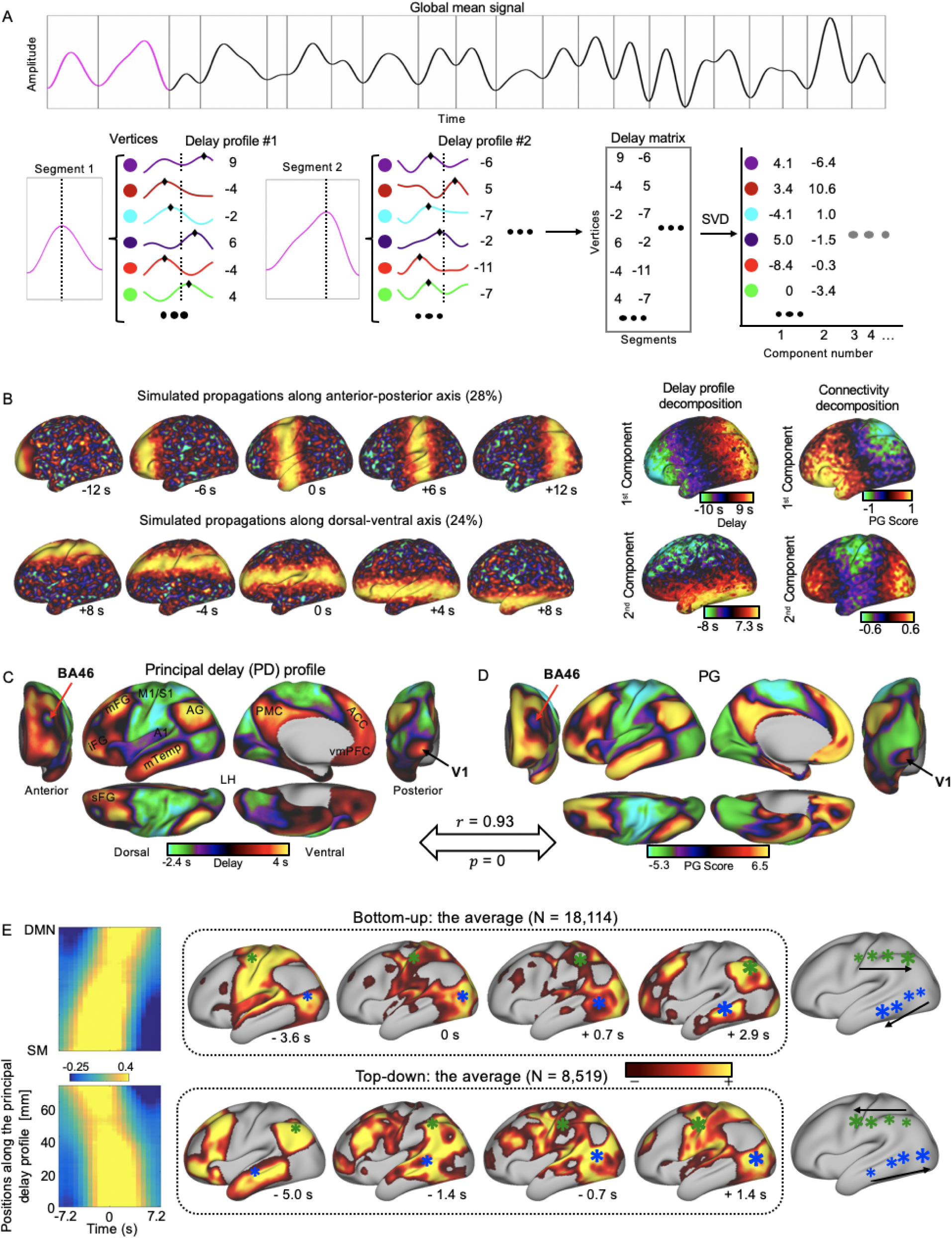
The major propagating direction of rsfMRI signals is highly similar to the principal gradient of rsfMRI connectivity. (A) Illustration of the procedures for deriving the principal delay profiles that represent the major propagating directions of rsfMRI signals. Specifically, the fMRI signals were cut into time segments based on the troughs of the global mean signal denoted by the gray vertical lines. Next, a delay profile was computed for each segment as the relative time delays of the local peak (black diamonds) at each cortical vertex with respect to the global peak (black dashed line). Finally, a singular value decomposition was applied to the delay matrix composed by delay profiles to extract the principal delay profile. (B) Decomposing delay profiles of synthesized fMRI data (left) using the proposed method recovered the directions of simulated propagating structures (middle). The decomposition of the connectivity matrix of the synthesized signals using the principal gradient method recovered the dominant direction but not the second one (right). (C) The application of the proposed method to real rsfMRI data produced the principal delay profile representing the principal direction of infra-slow rsfMRI propagations, which is extremely similar (*r* = 0.93, *p* = 0) to (D) the principal gradient of rsfMRI connectivity, including detailed features at the primary visual cortex (V1) and Brodmann area 46 (BA 46). (E) The averaged bottom-up (top, *N* = 18114) and top-down (bottom, *N* = 8519) propagations as presented on the time-position graphs (left) and the brain surface (right). Abbreviation: iFG, inferior frontal gyrus; mFG, middle frontal gyrus; AG, angular gyrus; mTemp, middle temporal cortex; M1, primary motor cortex; S1, primary somatosensory cortex; A1, primary auditory cortex; sFG, superior frontal gyrus; PMC, posteromedial cortex; ACC, anterior cingulate cortex; vmPFC, ventromedial prefrontal cotex; V1, primary visual cortex; BA 46, Brodmann area 46.

Decomposing the delay profiles of the rsfMRI segments of global involvement generated a principal delay profile (**Fig. 2C**), i.e., the first principal component (4.22% of the total variance, see **Fig. S3** for additional components), that is extremely similar (*r* = 0.93 across 59,412 vertices, *p* = 0) to the principal gradient of rsfMRI connectivity (**Fig. 2D**). In particular, the primary visual cortex (V1) appears to be an outlier for the overall hierarchical arrangements suggested by both maps. This deviation is however consistent with our observation that the V1 co-activates with the default mode network in the cross-hierarchy propagations (**Fig. 1C** for a single instance example). This principal delay profile is highly reproducible across sessions and subject groups (**Fig. S4**) and distinct from the lag map measured through dynamic susceptibility contrast MRI scans (*r* = 0.0035, *p* = 0.39; **Fig. S5**) (26). We then projected the rsfMRI signals onto this principal propagating direction and identified segments showing a significant (*p* < 0.05, compared with a pooled null distribution from the four control directions) time-position correlation. The top-down propagations (*N* = 8,519) and bottom-up propagations (*N* = 18,114) were found to account for 9.08% and 19.7% of the total time respectively with an average speed of 13.45 ± 7.78 mm/sec and 13.74 ± 7.51 mm/sec (mean ± SD) respectively. We then obtained the averaged patterns of these two types of propagations in both the time-position graph and brain surface (**Fig. 2E**), which are consistent with those of single instances (**Fig. 1C**). The above analyses were repeated on the rsfMRI signals going through different spatial and temporal filtering procedures, and the major findings about the cross-hierarchy contrast in the principal delay profile remained highly similar (**Fig. S6**). Thus, the infra-slow rsfMRI propagations are primarily along a hierarchical axis linking the higher-order and the lower-order brain regions, as represented either by the principal gradient or our principal delay profile.

### Infra-slow propagations in monkey ECoG signals follow a similar cross-hierarchy trajectory

To determine whether similar propagations are present in electrophysiological data free of hemodynamic contributions, we applied the same method to large-scale ECoG recordings from 4 monkeys in an eyes-closed rest condition. Using the same dataset, we have previously identified brain networks highly similar to resting-state connectivity networks based on power signal correlations (33). We first focused on the gamma-band (42–95 Hz) power that is known to be tightly linked to fMRI signals (34). The principal propagating direction (detected as the second component in one of 4 monkey) obtained by decomposing delay profiles of ECoG gamma-power showed a clear cross-hierarchy contrast between the sensory/motor areas and the higher-order frontal, anterior temporal, and parietal regions (**Fig. 3A**). This pattern, which is reproducible in all 4 monkeys (**Fig. 3A** and **Fig. S7**), is inversely similar (*r* = −0.71 ± 0.067, *p* < 10^−16^, see **Fig. S8** for electrode mapping on a macaque brain surface (35)) to the cortical myelination map that has been suggested to be a good approximation of cortical hierarchy (36). To further validate the existence of infra-slow propagations, we projected the ECoG gamma power signals onto this propagating direction in a similar way as the human rsfMRI analysis. The resulting time-position graphs clearly contained tilted bands with significant time-position correlations of local peaks, which are corresponding to the cross-hierarchy propagating activity on the brain surface (**Fig. 3B** and **Animations 3-4**). The time-position correlations for this cross-hierarchy axis showed a heavy tailed distribution that is significantly different (*p* = 1.65×10^−34^, two-sample Kolmogorov-Smirnov test) from that of control directions (**Fig. 3C**), which were obtained by rotating the principal delay profiles at a random angle to preserve the spatial continuity of electrodes (**Fig. S7**). Similar to the human results, the distribution is also asymmetric and characterized by a much larger peak for positive time-position correlations that suggest more bottom-up propagations. We then identified ECoG segments with propagating activity based on the time-position correlations and averaged them to obtain the mean propagating patterns of ECoG gamma power (**Fig. 3D**), which are similar to those of single instances.

**Fig. 3.**
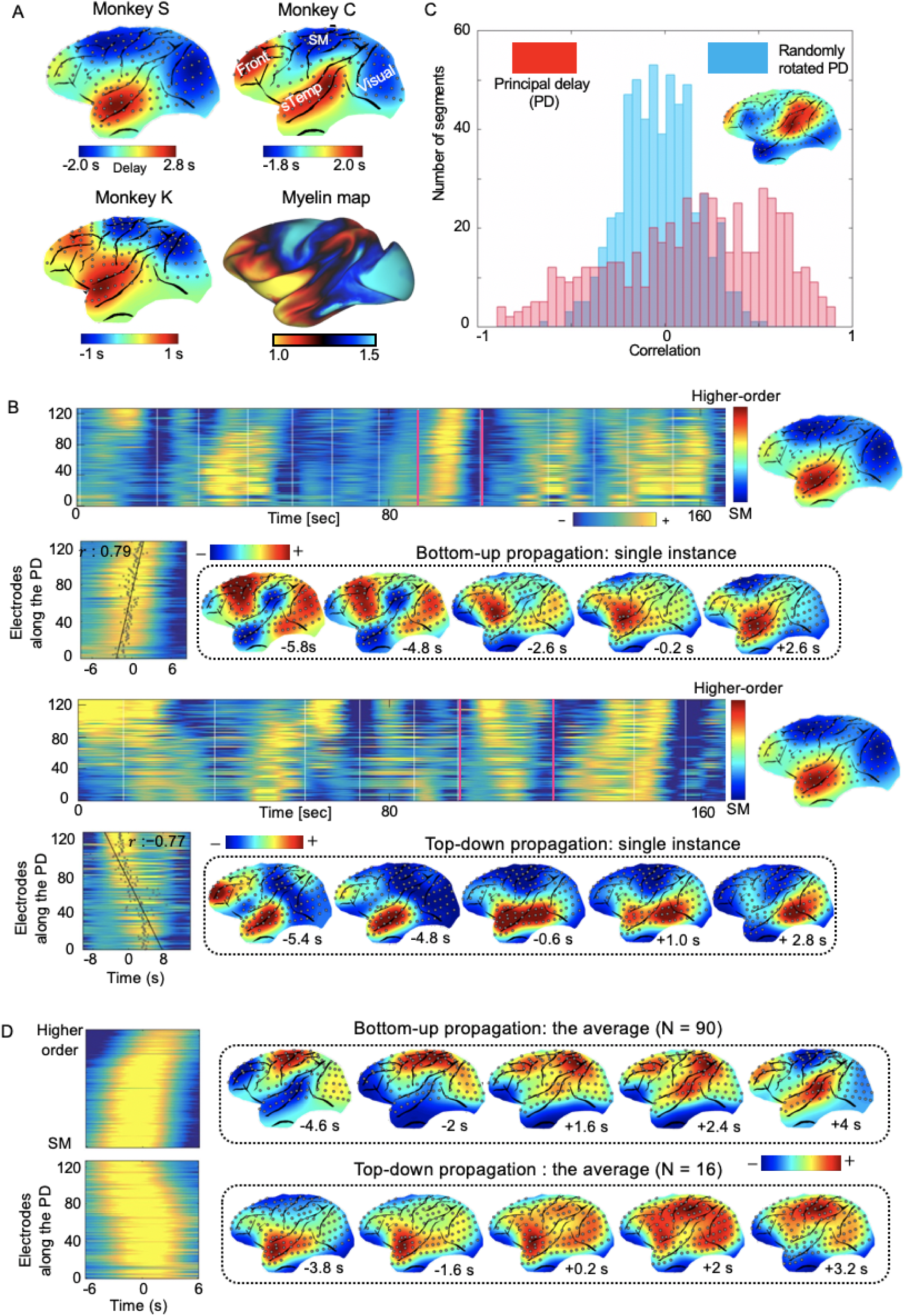
Cross-hierarchy propagations in the monkey ECoG signals. (A) The principal delay profile of the ECoG gamma-band (42–95 Hz) power shows a clear contrast between the sensory/motor (SM) areas and the higher-order (HO) brain regions, including the frontal, anterior temporal, and parietal cortices. This pattern is consistent across monkeys and inversely similar to the cortical myelination map of the monkey, which has been suggested as a good surrogate for estimating the cortical hierarchy (36). (B) Single exemplary instances of the top-down (i.e., from the higher-order to the sensory/motor regions) and bottom-up (i.e., from the sensory/motor to the higher-order regions) propagations as shown in the time-position graphs and on the brain surface. (C) The time-position correlations of the ECoG gamma-power segments for the principal propagating direction show a heavy tailed distribution that is significantly different (*p* = 1.65×10^−34^, two-sample Kolmogorov-Smirnov test) from the one obtained for control directions. The distributions were obtained by pooling the results from all four monkeys. (D) The averaged patterns of the top-down (*N* = 16) propagations for monkey S and bottom-up (*N* = 90) propagations for monkey C as shown in the time-position graphs (left) and the brain surface (right). Abbreviation: SM, sensory/motor region; sTemp, superior temporal cortex; Front, frontal cortex; Visual, visual cortex.

To know whether similar cross-hierarchy propagations are also present in resting-state brain activity of other frequency ranges, we repeated the same analysis for power signals of four other bands, i.e., delta (1–4 Hz), theta (5–8 Hz), alpha (9–15 Hz), and beta (17–32 Hz) bands, as well as the infra-slow (<0.1 Hz) band of the raw signals. It appeared that the principal delay profiles for the powers of the lower frequency bands are more characterized by a big contrast between the somatosensory/motor areas and the visual regions (**Fig. 4A**), and their spatial correlation with the cortical myelination map of monkeys is significantly (*p* < 0.01 for all four bands compared with the gamma band) lower than the gamma-band power (**Fig. 4B** and **Fig. S9-13**). Likewise, the principal delay profile of the infra-slow ECoG signals also had a significantly lower (*p* = 0.0135) correlation with the cortical myelination map than the gamma-band power without showing a clear cross-hierarchy contrast (**Fig. 4B**). Since it has been suggested that the low (30-80 Hz) and high (80-150 Hz) gamma activity may originate from different sources (37), we derived the principal delay profiles separately for the powers of the low and high gamma bands. Similar infra-slow propagations are present in the powers of these two gamma bands (**Fig. S14**). We repeated the same analyses on the ECoG gamma-band power going through different spatial and temporal filtering procedures, and the cross-hierarchy contrast in the principal delay profile can also be seen (**Fig. S15**). In summary, the gamma-band power of monkey ECoG signals exhibited infra-slow propagations similar to human rsfMRI in terms of the time scale (∼5-10 seconds) and, more importantly, the dominant propagating direction across the cortical hierarchy.

**Fig. 4.**
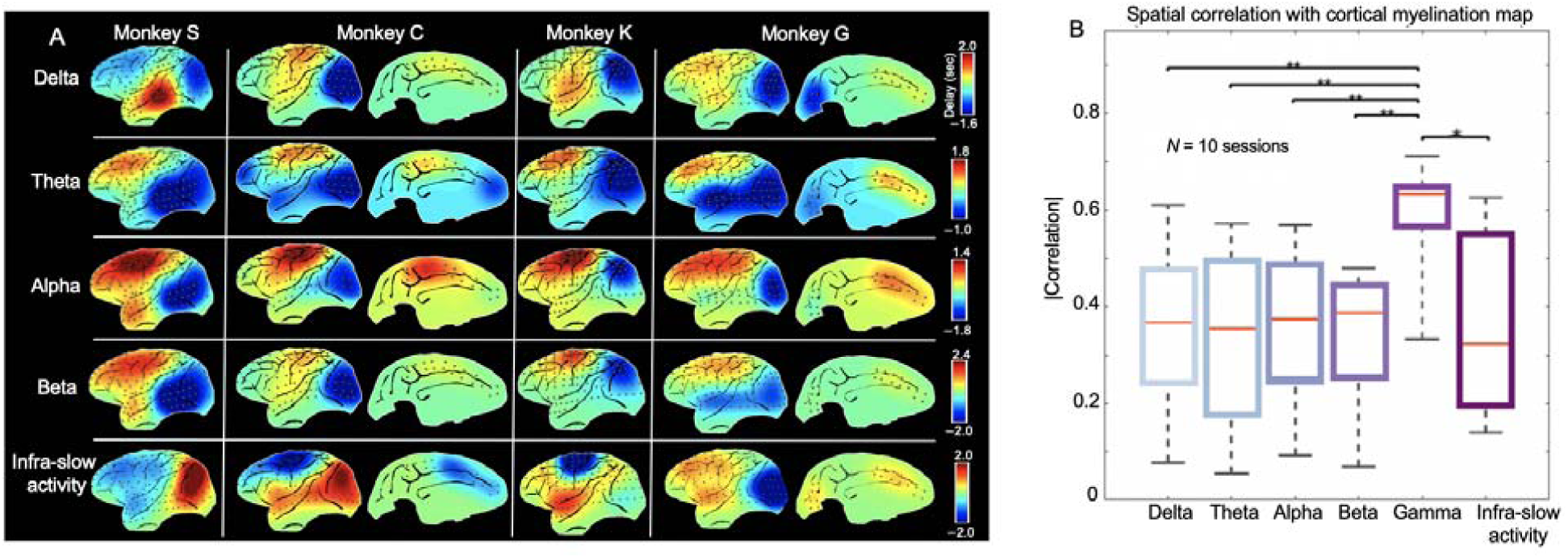
A reproducible cross-hierarchy propagating direction was present most strongly in the gamma-band power of ECoG signals. (A) The principal delay profiles derived for the ECoG powers of four other frequency bands: delta, 1–4 Hz; theta, 5–8 Hz; alpha, 9–15 Hz; beta, 17–32 Hz, as well as the infra-slow band (< 0.1 Hz) of ECoG signals (not power). (B) The spatial similarity between different principal delay profiles and the monkey cortical myelination map that has been suggested to be a good approximation of cortical hierarchy (36). The principal delay profiles and associated statistics were derived with respect to ten 25-minute eyes-closed sessions from all the monkeys. The error bar represents the standard error of mean. Asterisks represent the level of significance: **: 0.001 *p* 0.01. *: 0.01 *p* 0.05.

### Fine-scale propagations within sensory modalities are against the direction of global propagations

The anomalous position of the V1 in the PG and the principal delay profile (**Fig. 2C**), i.e., having similar scores/delays with the DMN, motivated us to examine fine-scale propagations within sensory modalities that are embedded in the global cross-hierarchy propagations. Within the visual system, the principal delay profile showed the most negative delay values at three isolated brain areas, including the MT+ complex, the dorsal stream visual cortex, and the ventral stream visual cortex (38), and increased its value towards the early visual cortex, as well as retinotopically from the periphery towards foveal areas (**Fig. 5A**). A simple linear regression confirmed a significant relationship between the delay value and the hierarchy level of brain regions (*p* = 3.3×10^−298^ for fV1-fV2-fV3-fV4-V4t-MT-MST-V6-V6A; and *p* = 0 for pV1-pV2-pV3-pV4-V4t-MT-MST-V6-V6A), which was defined based on a previous study (39) (**Fig. 5B**). This pattern is consistent with the trajectory of the bottom-up (i.e., sensory/motor to DMN) propagations within the visual cortex **(Fig. 5C**), which is actually from high-hierarchical visual areas to lower low-hierarchical ones. We then had closer inspection of the principal delay profile within the auditory and somatosensory systems to see whether the similar trend is present. Within the auditory system, the principal delay profile displayed a clear gradient across hierarchies from the A4, to belt regions, and then to A1 (*p* = 0) (**Fig. 5D** and **5E**), which is again consistent with the bottom-up propagation of local rsfMRI peaks (**Fig. 5F**). Local propagations within somatosensory system (**Fig. 5G** and **Fig. 5H**) appeared to follow more closely the somatotopic arrangement and show a strong contrast between limbs areas and eyes, face, and trunk areas (**Fig. 5I**). Nevertheless, the principal delay profile indeed showed a gradual and significant (*p* = 6.1×10^−313^) increase of value from the Brodmann area 2 (BA 2), to BA 1, and then to BA 3b and BA 3a (**Fig. 5J**). Similar analyses were also performed for the PG map, and the weaker but still significant relationships were found between the PG value and the hierarchical level (**Fig. 5** and **Fig. S16**). Outside the sensory systems, the very negative delays were also found in the frontal eye field (FEF), intraparietal sulcus (IPS), and BA 46 (**Fig. 5K**). Altogether, these regions compose a task-positive network known to have strong negative rsfMRI correlations with the default-mode network (40). To summarize, the local propagations within the sensory systems, which are embedded in the global cross-hierarchy propagations, appear to start/end at the sensory association areas and are opposite to the overall direction of the global propagation.

**Fig. 5.**
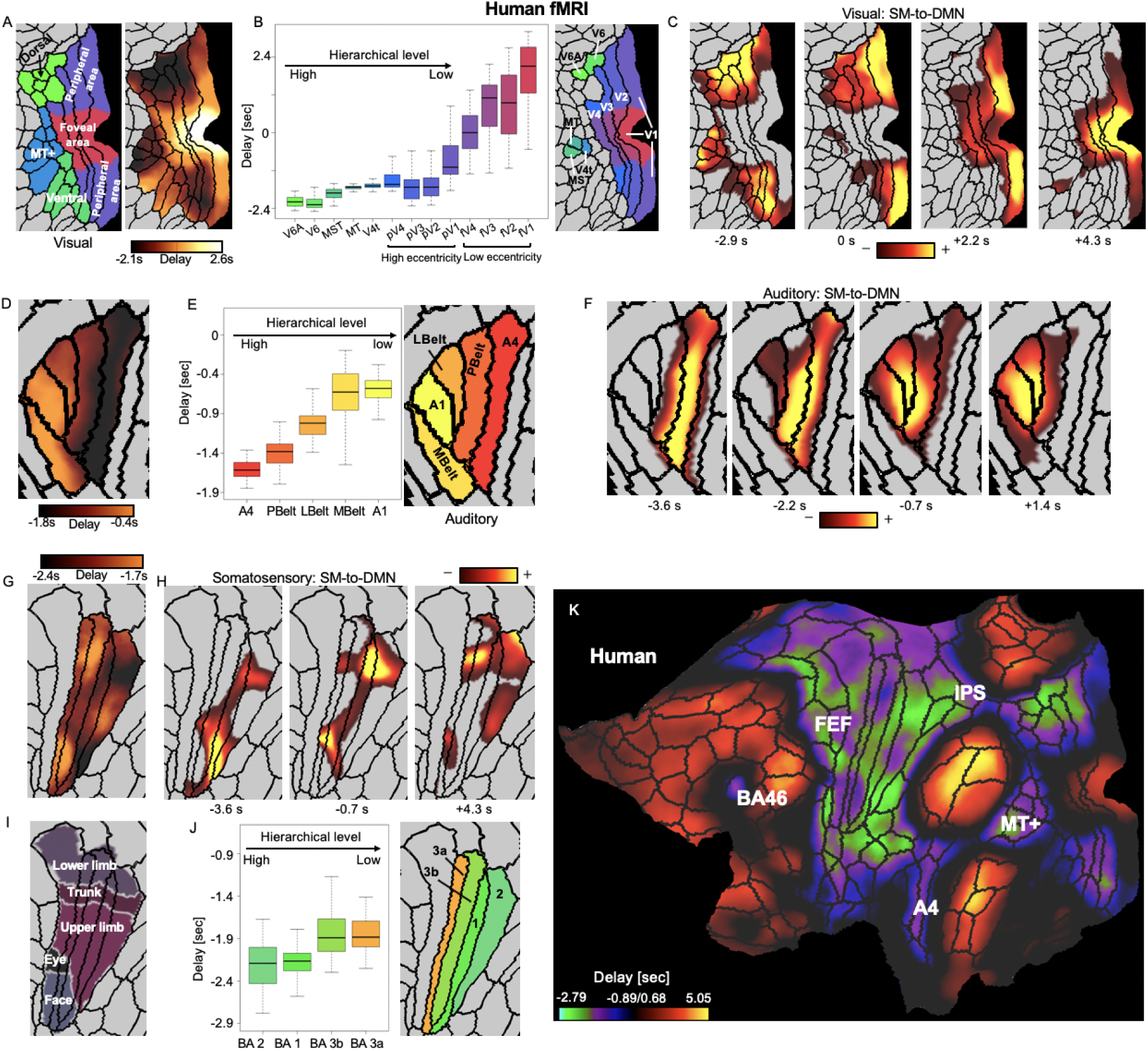
Local embedded propagations within the sensory modalities in human fMRI signals. (A) The local contrast of the principal delay profile within the three visual-related regions defined by a multi-modal parcellation atlas (38). (B) The averaged delay values of 13 visual parcels that were arranged according to their hierarchical (39) and retinotopic relationships (84). It should be noted that we do not assume any hierarchical relationship between the foveal and peripheral visual parcels. (C) The local trajectory of the bottom-up propagation within the visual system closely follows the principal delay profile, i.e., from the visual association areas to the early peripheral visual areas and then to the early foveal visual regions. (D-F) Results for the auditory system indicated a similar contrast and local propagation between the primary and association auditory areas. (G-I) Results for the somatosensory system show a weaker but still significant relationship between the delay and hierarchical level of 4 somatosensory parcels. The contrast of the principal delay profile also shows certain correspondence with (J) the somatotopic arrangement (85). (K) The principal delay profile on a flat brain surface suggests a few other brain regions outside the sensory systems showing large negative delays. Together with the sensory association areas, they compose the task-positive regions that have been shown previously to have strong negative-correlation with the DMN (40). Abbreviation: V6A, area V6A; V6, sixth visual area; MT+, MT+ complex; MST, medial superior temporal area; MT/V5, middle temporal area/fifth visual area; V4t, V4 transition zone; V4, fourth visual area (pV4 and fV4 are peripheral and foveal V4 respectively, same thereafter); V3, third visual area; V2, second visual area; V1, primary visual cortex; A4, auditory 4 complex; PBelt, parabelt complex; LBelt, lateral belt complex; MBelt, medial belt complex; A1, primary auditory cortex; BA, Brodmann’s area; FEF, frontal eye fields; IPS, intraarietal sulcus area; BA46, Brodmann area 46 SM, sensory/motor; DMN, default mode network.

### Subcortical co-activations/de-activations associated with the cross-hierarchy propagations

We then examined subcortical changes associated with these cortical propagations for additional evidence for its neural origin and also for important clues for underlying mechanisms. We averaged, in the volume space, the rsfMRI segments showing the propagations and converted them to Z-scores that represent the significance level of the deviation from the temporal mean. We found that the bottom-up cortical propagation is associated with strong, sequential co-activations/de-activations in specific subcortical regions. At the very early phase (*t* = −5.0 sec, with respect to the global signal peak) of this propagation, the weak co-activations at the sensory/motor regions are accompanied by strong de-activations in the default-mode network and extended areas. These cortical changes are associated with strong thalamic de-activations at the anterior nuclei (AN; peak Z: -24.41, mean Z: -12.98) and the dorsal part of the parvocellular division of the mediodorsal nucleus (MDpc; peak Z: -24.43, mean Z: -7.19), and to a less extent at the lateral dorsal (LD; peak Z: -19.56, mean Z: -10.73), the ventral lateral (VL; peak Z: -17.42, mean Z: -8.11), and the central lateral (CL; peak Z: -26.45, mean Z: -8.73) nuclei of the thalamus (**Fig. 6A**). In contrast, the significant thalamic co-activations are mostly confined at the anterior pulvinar (PuA; peak Z: 8.98, mean Z: 5.05). Starting from this time point, the thalamic co-activations started to spread from the PuA first to the posterior and ventral parts of the thalamus, which include many sensory relay nuclei, such as the ventral posterior medial (VPM), the ventral posterior lateral (VPL) nuclei and other parts of the pulvinar, and later to the AN and MDpc (**Fig. 6B-D**). Along this process, the cortical and thalamic co(de)-activations show striking correspondence consistent with known anatomical connections. For example, the V1 de-activation (*t* = −5.0 sec) is associated with very specific de-activation at the lateral geniculate nucleus (LGN) (**Fig. 6A**, the first row), and the maximal A1 co-activation (*t* = 1.4 sec) is also accompanied by the peak co-activation at the medial geniculate nucleus (MGN) (**Fig. 6C**, left). Outside the thalamus, a number of brainstem regions, including multiple nuclei related to arousal regulation, i.e., the dorsal raphé (DR; peak Z: -13.17, mean Z: -6.73), the median raphé (MR; peak Z: -9.07, mean Z: -6.14), the pendunculopontine nucleus (PPN; peak Z: -8.46, mean Z: - 5.65), the ventral tegmental area (VTA; peak Z: -11.02, mean Z: -6.79), and the locus coeruleus (LC; peak Z: -9.46, mean Z: -6.23) showed significant de-activations at very early phase (*t* = −6.5 sec) of the bottom-up propagation (**Fig. 6E**), along with the strong cortical de-activations at the precuneus and the cingulate (**Fig. 6E**, top right). The early de-activation of these brainstem nuclei was followed by a slow and gradual de-activation of three subcortical regions of arousal relevance, including the nucleus basalis (NB), the ventral part of the nucleus accumbens (NAcc) at the basal forebrain, and the substantia nigra (SN), which reached their peak de-activations around the middle of this propagation (*t* = 0) with widespread cortical co-activations (**Fig. 6F** and **6G**). Interestingly, the subcortical dynamics at the top-down propagation does not simply mirror that of the bottom-up propagation (**Fig. 6** and **Fig. S17**). Most notably, the strong de-activations are largely absent for the AN/MD and the brainstem nuclei throughout the top-down propagation (**Fig. 6** and **Fig. S17**). In summary, the cortical propagations are associated with co-activations and de-activations in corresponding thalamic nuclei, as well as the de-activation of subcortical regions of arousal relevance.

**Fig. 6.**
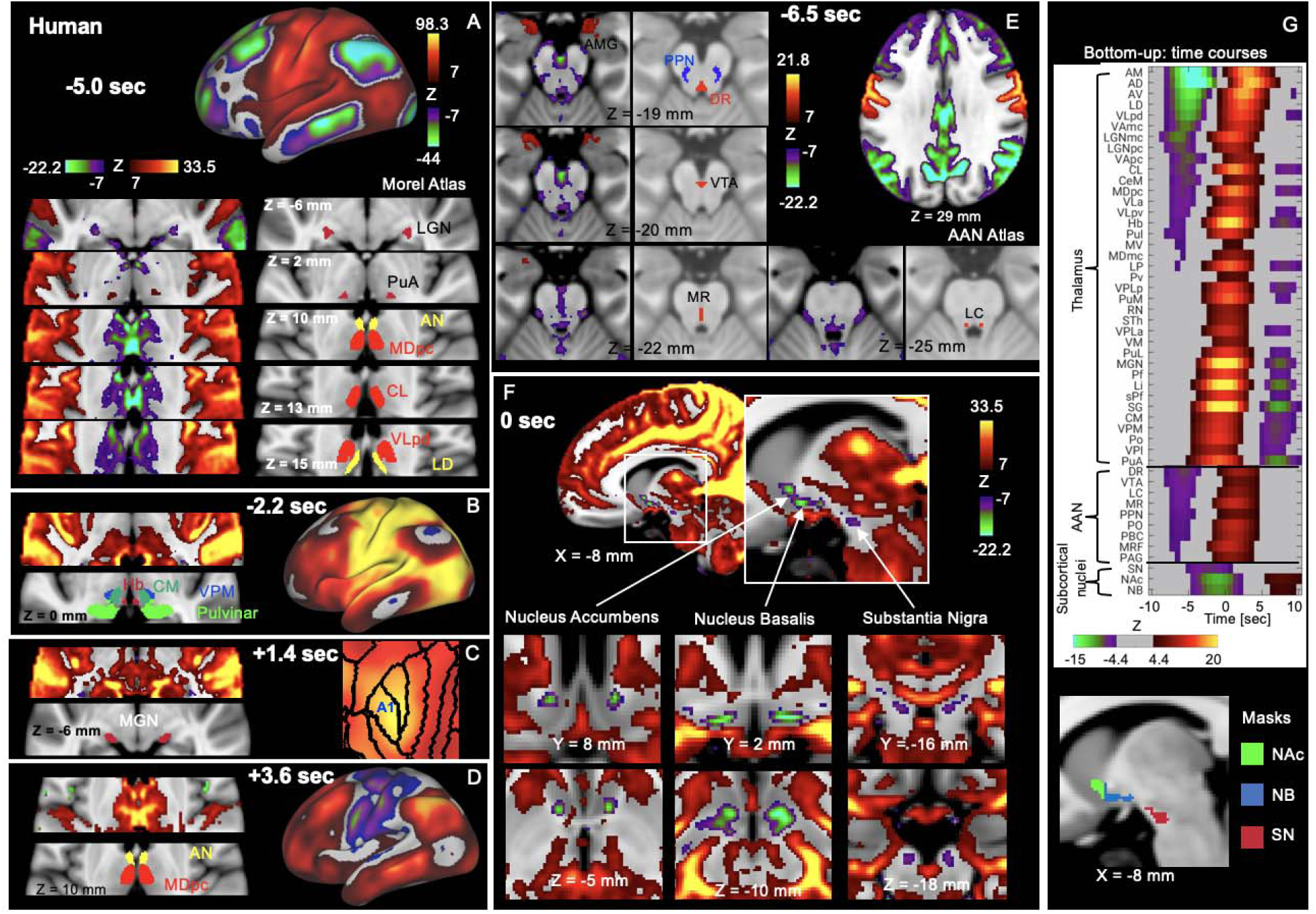
Subcortical co-(de)activations associated with the bottom-up propagation. (A-D) The thalamic co-activations/de-activation at different phases of the bottom-up propagations show a good correspondence with the cortical changes. The thalamic nuclei were located using the Morel Atlas (86). The time is with respect to the global mean peak. (A) The early de-activation of the default-mode network and V1 are associated with the thalamic de-activations in a few higher-order nuclei, particularly the AN and a part of MDpc, as well as the LGN, but the thalamic co-activation is only limited to the PuA. The thalamic co-activations then spread first to the posterior and ventral part of the thalamus (B) and eventually to the AN and MDpc at very late phase (D). (C) The MGN shows specific co-activations with the maximal A1 co-activation. (E) The earliest phase of the bottom-up propagation (*t* = −6.5 sec) involves the de-activations of a few brainstem nuclei of the ascending arousal system (AAS), which were located using the Harvard Ascending Arousal Network Atlas (87). The plots also showed the early co-activation of the AMG in the brainstem. (F) Following the early brainstem de-activations, the nucleus accumbens (NAc) at the ventral striatum, the nucleus basalis (NB) at the basal forebrain, and the substantia nigra (SN) at the brain stem started to slowly de-activate and reach the plateau with the widespread cortical co-activation. (G) The temporal dynamics of the subcortical regions shown in (A-F). The Z-score time courses were averaged within 37 thalamic regions of interest (ROIs) defined by the Morel’s atlas and 9 brainstem ROIs defined by the Harvard AAS atlas, as well as the three ROIs (NAc, NB, and SN) we defined by combining our results with corresponding brain atlases (80) (see the bottom for the masks we used). Each group of ROIs were sorted according to their value at *t* = −5.8 sec. The q-value after FDR corresponding to z-score 4.4 and 7 is 10^−5^ and 7.4×10^−13^ respectively. Error bars represent the standard error of the mean (SEM). Asterisks represent the level of significance: *: 0.01 *p* 0.05; **: 0.001 *p* 0.01; ***: *p* 0.001. Abbreviation: AMG, amygdala; MDmc, mediodorsal nucleus magnocellular part; MDpc, mediodorsal nucleus parvocellular part; MV, medioventral nucleus; CL, central lateral nucleus; CeM, central median nucleus; CM, centre median nucleus; Pv, paraventricular nucleus; Hb, Habenular nucleus; Pf, parafascicular nucleus; sPf, subparafascicular nucleus; PuM, medial pulvinar; PuI, inferior pulvinar; PuL, lateral pulvinar; PuA, anterior pulvinar; LP, lateral posterior nucleus; MGN, medial geniculate nucleus; SG, suprageniculate nucleus; Li, limitans nucleus; Po, posterior nucleus; LGN, lateral geniculate nucleus; VPLa, ventral posterior lateral nucleus anterior part; VPLp, ventral posterior lateral nucleus posterior part; VPM, ventral posterior medial nucleus; VPI, ventral posterior inferior nucleus; VLa, ventral lateral anterior nucleus; VLpd, ventral lateral posterior nucleus dorsal part; VLpv, ventral lateral posterior nucleus ventral part; VAmc, ventral anterior nucleus magnocellular part; VApc, ventral anterior nucleus parvocellular part; VM, ventral medial nucleus; AD, anterior dorsal nucleus; AM, anterior medial nucleus; AV, anterior ventral nucleus; LD, lateral dorsal nucleus; AN, anterior nucleus; STh, subthalamic nucleus; DR, dorsal raphe; VTA, ventral tegmental area; LC, locus coeruleus; MR, median raphe; PPN, pedunculopontine nucleus; PO, pontis oralis; PBC, parabrachial complex; MRF, midbrain reticular formation; PAG, periaqueductal gray; NAc, nucleus accumbens; NB, nucleus Basalis; SN, substantia Nigra.

### The modulation of the cross-hierarchy propagations across brain states of vigilance

The de-activations of arousal-related subcortical regions suggested a potential link between the bottom-up propagation and the brain arousal. Consistent with this notion, the rsfMRI lags between different brain regions were found to be completely reversed from wake to sleep in humans (41), and also from wake to anesthesia in mice (18). We thus suspected that the cross-hierarchy propagations in two opposite directions are sensitive to changes of brain arousal state. To test this hypothesis, we divided all rsfMRI sessions into three groups with the low, medium, and high level of arousal according to an adapted fMRI-based arousal estimation (42) and then compared their cross-hierarchy propagations. The ratio of the cross-hierarchy propagations in the two opposite directions is significantly different (*p* = 0 high vs. medium; *p* = 0 high vs. low; permutation test) in the three groups with the high arousal group having relatively less bottom-up propagations (**Fig. 7A** and **Fig. S18**). The fMRI-based arousal estimation involved only a template-matching process that is not expected to introduce any bias towards any propagating directions. To have a more independent estimation of brain arousal level, we also computed this ratio in a subset of the sessions in which subjects were noted by experimenters to be sleeping during the rsfMRI scanning. This subset of rsfMRI sessions showed significantly lower (*p* = 0.021, permutation test) ratio compared with other sessions (**Fig. 7B**). A similar comparison was made for the cross-hierarchy propagations in monkey ECoG gamma powers across three experimental conditions: a more alert eyes-open condition, a more sleep-conducive eyes-closed condition, and the sleep condition. Consistent with the human rsfMRI results, the ratio of the top-down propagations to the bottom-up propagations are significantly different (*p* = 0.0233 eye-open vs. eye-closed; *p* = 0.0045 eye-open vs. sleep; *p* = 0.029 eye-closed vs. sleep; permutation test) across the three conditions with the sleep state showing much more bottom-up propagations and less top-down ones (**Fig. S18**). To summarize, both monkey electrophysiology and human rsfMRI data suggest that the state of lower arousal is associated with less top-down but more bottom-up propagating activity.

**Fig. 7.**
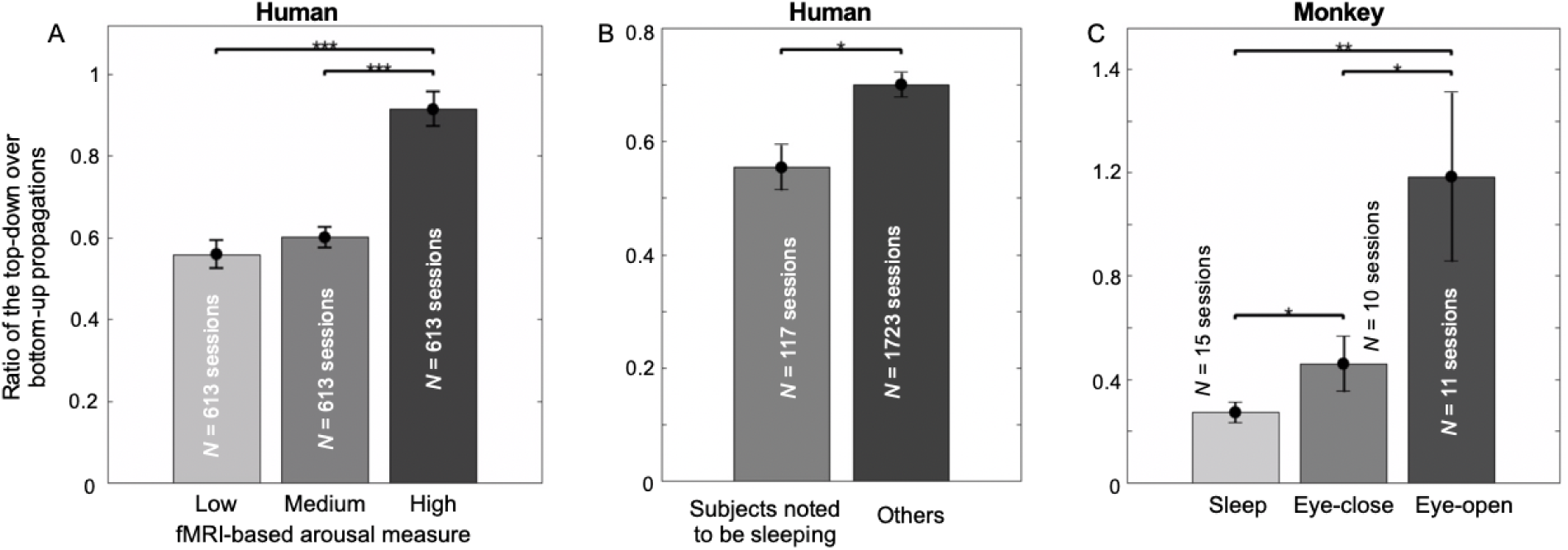
The cross-hierarchy propagating activity is sensitive to brain arousal level. (A) The ratio of the top-down propagation to the bottom-up propagation is significantly different in three groups of sessions showing distinct arousal levels as measured by an fMRI-based arousal index (89). (B) The same ratio is also significantly lower in a subset of sessions in which subjects were noted to be sleeping during rsfMRI scanning. (C) The ratio of the top-down propagation to the bottom-up propagation in the monkey ECoG gamma powers shows a similar and significant modulation across the eyes-open, eyes-closed, and sleep sessions. Error bars represent the standard error of the mean (SEM). Asterisks represent the level of significance: *: 0.01 < *p* ≤ 0.05; **: 0.001 < *p* ≤ 0.01; ***: *p* ≤ 0.001.

## Discussion

Here we showed that the resting-state brain activity, measured by fMRI in humans or electrophysiology in monkeys, is characterized by distinctive propagations sweeping the cortex in two opposite directions along an axis. This trajectory is extremely similar to, and thus these spatiotemporal propagations may underlie, the principal gradient of rsfMRI connectivity (31). The cross-hierarchy ECoG propagations are present most strongly in the gamma-band power. The local propagations within the sensory modalities are in a direction opposite to the overall direction of the global propagation, suggesting that these neuronal processes start/end at the sensory association regions. The bottom-up propagation is associated with sequential co-(de)activations at specific subcortical nuclei, including many related to arousal regulation. Consistent with this finding, the cross-hierarchy propagations are significantly modulated by the brain arousal level. Overall, the findings from this study supported the neural origin of the infra-slow rsfMRI propagations, revealed detailed features and behavioral relevance of infra-slow propagating activity, and also linked it to the principal gradient of rsfMRI connectivity.

The study added direct evidence for the neural origin of rsfMRI propagations by showing corresponding electrophysiological propagations on a similar time scale, along a similar direction, and showing a similar state-dependency. Inferring the propagating activity with fMRI signals could be problematic given the known region-specific hemodynamic delays (21). This concern became heightened given a series of studies showing that the systematic low-frequency oscillations of blood signals cause rsfMRI signal delays consistent with the blood transition time (22–26). Some indirect evidence has been used to argue against the vascular origin of the infra-slow rsfMRI propagations. For example, the rsfMRI lag structures/threads persist after regressing out the vascular time lags (43) and are sensitive to brain state, which are not expected from a vascular based propagation (18, 41). Consistent with the previous findings, we found no similarity (*r* = 0.0035 and *p* = 0.39, **Fig. S5**) between the principal delay profile and the vascular lags as measured by the dynamic susceptibility contrast MRI (26), as well as a strong dependency of the cross-hierarchy propagations on brain state (**Fig. 7**). Moreover, the detailed features of the cross-hierarchy propagations, including the existence of propagating instances in opposite directions (**Fig. 2**), their fine-scale trajectories within the sensory systems (**Fig. 5**), the sequential involvement of specific subcortical nuclei (**Fig. 6**), good correspondences between cortical and thalamic co-activations/de-activations (**Fig. 6**) added further evidence against their vascular origin.

The study provided additional details about the infra-slow propagating activity. First, the cross-hierarchy propagations are present much more strongly in the gamma-band power than other bandlimited powers and the infra-slow band of the raw signals. This explains the previous finding that the long-range ECoG power correlations between the high-order regions, can only be found in the gamma band (33). Given that the gamma activity is correlated with neuronal firing rates (44), the cross-hierarchy propagations of gamma-band power may represent an excursion of cortical excitation sweeping along the hierarchical axis. Although it has been suggested that the evoked low (30-80 Hz) and high (80-150 Hz) gamma powers originate from different sources (37), we observed similar cross-hierarchy propagations in these two sub-gamma-bands. Either the spontaneous gamma-band activity may not contain two separate components, or they both contain similar cross-hierarchy propagations. The infra-slow (<0.1 Hz) ECoG signals (not the power) have been shown to have structured correlations similar to rsfMRI (45), but we failed to identify similar cross-hierarchy propagations from them (**Fig. 4**). However, we cannot exclude the possibility that the monkey ECoG was not optimized for and thus fail to record the infra-slow activity (46). Secondly, the fine-scale spatiotemporal dynamics were also elucidated within the sensory modalities. Specifically, the cross-hierarchy propagations appear to start/end at the unimodal association areas, rather than the primary areas. Thirdly, the cross-hierarchy propagating activity is conserved across human and monkeys. The fast (< 1 sec) propagating brain activity has also been studied in mice mostly using the optical imaging, but often described as along the anterior-posterior axis (27, 29). Close inspection has suggested that they may indeed follow more specific trajectories between the sensory/motor regions and higher-order cortices (18, 28), and it remains to be determined whether the infra-slow propagating activity in rodents is also aligned with the cortical hierarchical gradient.

The specific subcortical co-activations/de-activations point to the involvement of the ascending arousal system in the bottom-up propagation, which might represent a brain process associated with transit arousal modulations. The earliest co-activations at the sensory association areas are accompanied by significant de-activations in multiple brainstem nuclei of the ascending arousal system, including the locus coeruleus, median raphe, pedunculopontine nucleus, and ventral tegmental area. The early brainstem de-activation is then followed by a slow and gradual de-activation of three other subcortical regions of arousal relevance, i.e., the substantia nigra, nucleus basalis, and nucleus accumbens (47). Importantly, the brainstem de-activation is also accompanied, if not triggered, by strong de-activations in a set of cortical and thalamic regions, including the default-mode network, the frontoparietal network, the precuneus, and the anterior and mediodorsal thalamus (**Fig. 6E**). This exact set of brain regions showed significant reduction in glucose metabolic rate and cerebral blood flow during anesthetic-induced unconsciousness (48), suggesting that the bottom-up propagating activity is associated with transient arousal modulations. Consistent with this hypothesis, its occurrence rate is significantly modulated across brain states of distinct arousal levels (**Fig. 7**). The large global rsfMRI peaks have been linked to a neurophysiological event indicative of arousal modulations (6), which may represent an instantaneous phase (at *t* = 0) of this bottom-up propagation, given their similar sensory-dominant cortical co-activations and subcortical de-activations (6).

While the subcortical ascending arousal system may be involved in the initiation of the cross-hierarchy propagations, the network mechanism underlying its cortical propagation remains elusive. The cross-hierarchy propagations are unlikely mediated through axonal conduction given its slow speed (∼5-25 mm/sec). This speed however fells into the velocity range (∼10-100 mm/sec) of spontaneous waves of depolarization observed in the barrel cortex of rodents through optical imaging and whole-cell recordings (49). It has been hypothesized that the recurrent excitation through local synaptic connections in layer 2/3 contributes to those spontaneous propagating waves. The similar mechanisms may also underlie the infra-slow propagations observed in this study. Moreover, the top-down and bottom-up propagations may take distinct routes cross cortical layers that are consistent with the known feedback and feedforward connections (50–52). The propagating speed is close to what has been observed for the propagation of epileptic activity (20-100 mm/sec) (53–56), but it remains unclear whether they share common mechanisms. Multimodal techniques capable of imaging brain activity across distinct spatial and temporal scales are required for a deep understanding of the mechanism underlying the cross-hierarchy propagations in the future.

Cross-hierarchy propagating activity may, in fact, underlies the principal gradient of rsfMRI connectivity (31). We showed an extremely high similarity (*r* = 0.93) between the principal propagating direction and the principal connectivity gradient, which is unlikely a coincidence. Instead, the propagations are expected to synchronize rsfMRI signals and affect their observed correlations in a systematic way along its trajectory. It is also worth noting that we failed to find a trajectory similar to the second motor-to-visual gradient of rsfMRI connectivity using the delay profile method (**Fig. S3**) or observe single instances of such propagation (**Fig. 1D**, the second column). Such a motor-to-visual contrast, which is also present in the vascular lag map (**Fig. S5**), may be caused by small but significant time delays observed between the motor and visual areas in the cross-hierarchy propagations. The cross-hierarchy propagating activity might also be related to other rsfMRI findings showing features related to the hierarchical axis. For example, converging evidence from rats, monkeys, and humans has suggested that rsfMRI connectivity/dynamics of the higher-order cognitive networks and lower-order sensory/motor networks are divergently modulated by anesthesia (57–60).

The overlap between the principal propagating direction and the hierarchical axis of the brain implies the functional significance of the infra-slow propagating activity. A speculation of its functional roles comes from its analogy to “propagations” in artificial neuronal networks (61). The learning of such large-scale, non-linear models requires efficient algorithms, which often involves iterative propagations of information across hierarchical layers, including a forward propagation of information and, more importantly, a backpropagation of model errors to optimize weights/connections in a successive manner (62). Such repetitive, sequential activations across hierarchical stages might be even more important for the modification of real neuronal synapses. The cross-hierarchy propagations would serve this purpose by creating successive excitations across the hierarchical axis of the brain. Consistent with this conjecture, the hippocampal ripples, a neuronal process tightly linked to learning and memory consolidation, have been found to be co-modulated with cortical delta-band power in a slow (∼0.1 Hz) rhythm comparable to the time scale of the infra-slow propagating activity (63), and also associated with massive fMRI cortical activations with region-specific delays, particularly at the V1, suggestive of propagating behavior (64).

## Materials and Methods

### HCP data and preprocessing

We used the human connectome project (HCP) 500-subject data release, including 526 healthy subjects scanned on a 3T customized Siemens Skyra scanner. We limited our analyses to 460 subjects (age: 22-35 years, 271 females) who completed all four 15-min resting-state fMRI (rsfMRI) sessions on two separate days (two sessions per day). The data were collected using multiband echo-planar imaging with an acceleration factor of 8 (65). The temporal and spatial resolution of the data are 0.72 sec and 2-mm isotropic respectively. Four 15-min scanning sessions of 460 subjects were used in our analysis.

The rsfMRI data were preprocessed based on (66) using FSL (67), FreeSurfer (68), and HCP workbench (69) and the HCP FIX-ICA denoising pipeline was applied to remove artifacts. The multimodal surface matching registration was used in the HCP dataset to improve inter-subject registration (70, 71). We used both rsfMRI surface and volume data. The rsfMRI cortical surface data were represented in standard HCP fs_LR 32k surface mesh and each hemisphere included 32,492 nodes (59,412 total excluding the non-cortical medial wall). We smoothed rsfMRI data both spatially on the fs_LR 32k surface using a Gaussian smoothing kernel (sigma = 2 mm) and temporally using bandpass filtering at 0.001 - 0.1 Hz, and then standardized each vertex’s signal by subtracting the mean and dividing by the standard deviation. For the rsfMRI volume data including both cortical and subcortical areas, we smoothed rsfMRI volume data temporally (0.001-0.1 Hz) and standardized each voxel’s signal by subtracting the mean and dividing by the standard deviation.

### ECoG data and preprocessing

The monkey electrophysiology dataset was downloaded from the website (http://neurotycho.org) and had been described in a previous publication (33). To sum up, all procedures were approved by the RIKEN ethics committee. An implanted customized 128-channel ECoG electrode array (Unique Medical, Japan) was used to record neural signals (72). Each ECoG electrode had a 3-mm diameter platinum disc with a 5 mm inter-electrode distance. We used the ECoG data from four adult macaque monkeys (monkey K, G, and C Macaca fuscata and monkey S Macaca mulatta). The 128-channel ECoG electrode array was implanted in the left hemisphere covering the majority of cortical regions. The reference and ground electrodes were implanted in the subdural space and the epidural space, respectively. ECoG recordings were conducted with a sampling rate of 1kHz using the Cerebus data acquisition system (Blackrock, UT, USA). More specific information can be found in (73).

The ECoG signals were recorded under three brain states: the eyes-open, eyes-closed, and sleep states. Under the eyes-closed waking state and the natural sleep state, the monkeys sat calmly in a dark and quiet environment with eyes covered. During the sleep condition, the slow-wave oscillations were observed intermittently on the ECoG data. Under the eye-open condition, the eye mask was removed. Experiments were conducted on separate days. The ECoG data under the eyes-open and eyes-closed conditions were available in all of the four monkeys. The ECoG data under the natural sleep condition were only available in monkey C and monkey G.

The total length of ECoG data under the eyes-closed condition was about 85 min, 89 min, 60 min, and 62 min for monkey C, monkey G, monkey K, and monkey S, respectively. The total length of ECoG data under the eyes-open condition was about 103 min, 85 min, 60 min, and 61 min for monkey C, monkey G, monkey K, and monkey S, respectively. The total length of ECoG data under the sleep condition was about 243 min and 157 min for monkey C and monkey G, respectively.

We removed the line noise at the primary frequency (50 Hz) and its harmonics using Chronux (74). We excluded three channels for monkey G, one channel for monkey K, and one channel for monkey S from subsequent analyses due to serious artifacts that cannot be removed. We re-referenced the ECoG signals to the mean of all channels. To extract the band-limited power signals, we first calculated spectrograms between 1 and 100 Hz using a multi-taper time-frequency transformation with a window length of 1 sec, a step of 0.2 sec and the number of tapers equal to 5 provided by Chronux (74). We then converted the power spectrogram into decibel units using the logarithmic function. Next, we normalized the power spectrogram at each frequency bin by subtracting the temporal mean and dividing by its temporal standard deviation. The normalized spectrogram was averaged within different frequency bands: delta 1 - 4 Hz; theta 5 - 8 Hz; alpha 9 - 15 Hz; beta 17 - 32 Hz; and gamma 42 - 95 Hz. The gamma frequency band was defined conservatively as 42–95 Hz, within which the power signals of different frequency bins show similar temporal dynamics. We also extracted the power of the low-(30-80 Hz) and high-gamma (80-150 Hz) bands as defined by a previous study (37). Then, the band limited power signals were smoothed both temporally using a low-pass filter (<0.1 Hz) and spatially using a Gaussian smoothing kernel (sigma = 5 mm), then standardized by removing the mean and dividing by its standard deviation. The effect of temporal filter used here was examined to make sure that it won’t produce any phase shifts (**Fig. S19**).

### Projecting the rsfMRI signals onto the principal gradient direction

The rsfMRI signals were projected onto the principal gradient (PG) (31) direction to generate time-position correlations as follows. The principal gradient was obtained by a previous study with applying the diffusion mapping, a low-dimensional embedding method, to a group averaged connectome matrix (31). First, we reduced the spatial dimension by sorting 59,412 cortical surface vertices according to the principal graident and then dividing these cortical surface vertices into 70 position bins of equal size. Next, the fMRI signals within each position bin were averaged to generate the time-position graph (see **Fig. 1A**). Secondly, the time-position graph was cut into time segments based on the troughs of the global mean signal. Thirdly, for each time segment, the Pearson’s correlation between the relative timing with respect to the global mean peak and position of local peaks across all the position bins was calculated. A strong positive time-position correlation would indicate a propagation of the fMRI signal along the principal gradient direction within this time segment and a strong negative time-position correlation would indicate a propagation opposite the PG direction. The local peak of each position bin was defined as the local maxima with a value larger than zero. If more than one local peak were detected, the local peak was defined as the one with the largest peak amplitude. The time-position correlation was only computed for time segments whose local peaks were identified in at least 56 (70 × 80%) position bins.

To focus our analysis on the time segments with a global involvement, we identified time segments with relatively large global peak amplitudes by using a threshold defined from a null distribution and calculated time-position correlations of those identified time segments. We first generated a null distribution of global peak amplitudes by randomly shifting the fMRI signals of position bins and then calculating the global peak amplitudes of those randomly shifted signals. The random shifts were uniformly distributed integers between 1 and the total number of time points (1200) in a single scanning session. The time segments with a global involvement were defined as those with a global peak amplitude exceeding the 99^th^ percentile of the null distribution, which account for 58.80 % of the total segments. We also calculated the delays between the global peak and the local peaks of 70 position bins for time segments of the real fMRI signals and randomly shifted signals.

For the time segments with a global involvement, the time-position correlations along four other control directions were also computed in the same way as described for the principal gradient. The four control directions included the second gradient of rsfMRI connectivity derived by the principal gradient study (31), and three artificial directions, i.e., anterior-to-posterior direction, dorsal-to-ventral direction, and randomly rotated principal gradient. The anterior-to-posterior direction map was generated by assigning increasing amplitude to vertices from posterior to anterior direction according to the y coordinate. The dorsal-to-ventral direction map was generated by assigning increasing amplitude to vertices from ventral to dorsal direction according to the z coordinate. The randomly rotated principal gradient was generated by rotating the principal gradient map on the spherical fs_LR 32k surface space with random degrees with respect to the x, y, z axes, which relocated and the principal gradient preserved the relative topology of the PG.

### Principal propagating direction in the human rsfMRI signals

A principal delay (PD) profile was derived by applying a singular value decomposition (SVD) (75) to delay profiles of time segments with a global involvement. For any given time segment with a global involvement, a delay profile was computed as the relative time delay of the local peak at each cortical surface vertex with respect to the global peak. The local peak of each vertex was defined as the local maxima with a value larger than zero. If more than one local peak were detected, which were rare, the local peak was defined as the one with the largest peak amplitude. We focused on the delay profiles with at least 47,530 (59,412×80%) local peaks. For those selected delay profiles, if the local peak amplitude was less than zero or no local maxima was detected, then its relative time delay was defined as the mean time delay of its three nearest vertices with local peaks larger than zero. Specifically, for each vertex without time delay, the distance on the brain surface between this vertex and other vertices with time delay was calculated and sorted. Then the averaged time delay of three vertices with smallest distance were used to replace the time delay of that vertex. The distance on the brain surface was calculated based on the coordinates of vertices. Next, a delay matrix was formed by concatenating all of the delay profiles, to which we then applied SVD to extract the principal delay profiles (see **Fig. 2A**). This delay profile decomposition method shares a similar idea with several previous approaches (76–78) in utilizing the time delays between brain regions to infer propagating brain activity.

SVD is to reduce high dimensional data to lower dimensions of uncorrelated components. The delay matrix is an *n*× *m* matrix, where n is the number of cortical vertices and *m* is the number of delay profile. The delay matrix was denoted as *X*. Applying SVD to *X* generates:

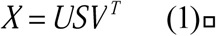

where *U* is an *n*× *n* orthonormal matrix, whose column represents left singular vector *S* is an n× m diagonal matrix, whose diagonal entries represent singular values of *X*. *V* is an *m*× *m* orthonormal matrix, whose column represents right singular vector. The columns of *U* were ordered based on the variance explained of *X*. The square of each diagonal singular value in *S* denoted the variance of the corresponding vectors. A principal delay profile is the first column of *U*, which explains the largest variance of *X*. It is expected to reflect the major propagating direction.

The rsfMRI time segments with a propagation along the principal delay profile were defined as described below. The rsfMRI segments with a global involvement were projected onto the principal delay profile direction and the corresponding time-position correlations were then calculated in the same way as described above. We repeated the same procedure for the four control directions describe above, and built a null distribution of time-position correlations by pooling time-position correlations for all the four control directions. A positive correlation exceeding the 1.64 standard deviation (SD) of the null distribution was regarded as a bottom-up (from sensory/motor regions (SM) to the default mode network (DMN)) propagation and a negative correlation exceeding 1.64SD of the null distribution was regarded as a top-down (from the DMN to SM) propagation. The 1.64SD was used because the critical value for a 90% confidence level with 5% on each side is 1.64. The total time of the top-down or bottom-up propagations was calculated as the total length of all the time segments identified to have a top-down or bottom-up propagation.

The SVD components captured the major propagating directions but in an arbitrary unit rather than in seconds. Therefore, we re-scaled the derived principal delay profile based on the time-position relationship of the time segments with top-down or bottom-up propagations. Specifically, we calculated the regression coefficient of the time-position relationship for each segment with a propagation to estimate the propagating speed with assuming the geodesic distance on the cortical surface between the primary sensory/motor regions and the default-mode network is 80 mm (31). We computed the averaged propagating speed across all the time segments with propagations and then utilized it to re-scale the principal delay profiles into the unit of seconds. Applying such rescaling procedure on the synthesized data successfully generated two principal delay profiles that are consistent with the duration of simulated propagating structures (**Fig. 2B** and **Fig. S2**).

The z-score maps for averaged bottom-up and top-down propagations were calculated in the surface and volume space as follows. First, the global peaks within the time segments with propagations were located. A time window of 15.12 sec (21TR × 0.72 sec/TR) was defined to contain ten time points before and ten time points after each of these global peaks, with setting the global peak point as time zero. Second, rsfMRI signals in both surface and volume space of these time windows were averaged respectively to obtain maps showing averaged propagation, which were then converted to z-score maps according to the formula below:

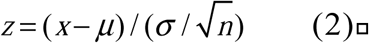

The normalized rsfMRI signals have a zero mean (µ = 0) and a unit standard deviation (a-= 1). n is the total number of time segments with propagations, and X is the averaged propagating map. The false discovery rate (FDR) is used to correct multiple comparisons. The function ‘fizt_t2p’ from AFNI (79) was used to covert the z-score to the p-value. The FDR q values were generated by using the 3dFDR program from AFNI (79). The associated time-position graphs of the defined time windows with propagations were also averaged.

The cerebrospinal fluid regions were masked out from the averaged top-down and bottom-up propagations in the volume space. The cerebrospinal fluid regions were defined based on the Harvard-Oxford subcortical structural probabilistic atlas (80) using a threshold of 30% probability.

We tested the reproducibility of the principal delay profile in humans using a split-half analysis. There are four sessions of rsfMRI data acquired in two different days. For each session of data, we randomly split subjects into two equal groups and calculated the principal delay profile for each group as described above.

The principal delay profile was also computed on fMRI signals with skipping the temporal filtering and/or spatial filtering, or switching their order (**Fig. S6**). For all these cases, the fMRI signals were also cut into time segments based on the global mean signal, calculated using the fMRIs signal filtered of filtered data. Otherwise, we would have time segments of a few seconds, which would stop any meaningful subsequent analyses.

### Simulation of rsfMRI signals with artificial propagations

To test whether the delay profile decomposition method and the principal gradient method can successfully identify the direction of propagating activity, we simulated fMRI signals on the brain cortical surface containing propagating structures along artificial directions and then applied these two methods to the simulated data. The propagating structures were generated by creating a spatial band of high amplitude signals and then shifting it along the anterior-to-posterior, posterior-to-anterior, or dorsal-to-ventral directions over time on the brain surface. The signal modulation within the activation band along its propagating direction was described as a Gaussian function. We simulated 6 types of propagating structures that propagated across the whole brain with two different speeds in three different directions, including those propagating through the whole brain in the anterior-posterior and the opposite directions in 19 and 29 seconds respectively, as well as those propagating across the whole brain in the dorsal-to-ventral directions in 11 and 20 seconds. The simulated propagating structures were convolved with a canonical hemodynamic response function from SPM (https://www.fil.ion.ucl.ac.uk/spm/) and then randomly inserted into background fMRI signals that were modeled as white noise. We randomly inserted two of each type of propagating structure along the anterior-to-posterior direction, one of each type of propagating structure along the posterior-to-anterior direction, three of each type of propagating structure along the dorsal-to-ventral direction into each session of simulated fMRI signals of 1200 time points. The temporal resolution of simulated fMRI signals was assigned as 1 sec arbitrarily. The simulated fMRI signals were further spatially smoothed using a Gaussian smoothing kernel (sigma = 2 mm) and temporally filtered using bandpass filtering at 0.001 - 0.1 Hz. We simulated a total of 50 sessions of fMRI data, and then derived their principal delay profile as described above and the principal gradient by decomposing the connectivity matrix using codes provided by the previous study (31). The procedure was repeated for two sets of simulated data with different signal-to-noise ratio (SNR), which were obtained by setting the peak signal of the activation bands as 2 and 5 times of the SD of background white noise.

### Principal propagating direction in the monkey ECoG data

The delay profile decomposition method described above was also applied to the ECoG data to derive the principal delay profile for different bandlimited powers, as well as the infra-slow (< 0.1 Hz) band of the raw signal (not power). Here we take the gamma-power as an example. The global mean signal was calculated as the averaged ECoG gamma-power across all electrodes. The signals were cut into time segments based on troughs of the global mean signal. Next, the time segments with a global involvement were identified if the global peak amplitude exceeded the threshold (the 99^th^ percentile of a null distribution of the global peak amplitudes). The null distribution was created by calculating the global peak amplitudes after randomly shifting the time series of each electrode. The random shifts were uniformly distributed integers between 1 and the number of the time points in the experimental session. The analyses below focused on the time segments with a global involvement.

A delay profile was generated for each time segment by computing the relative time delays between the local peak of individual electrodes and the global peak. The local peak of each electrode was defined as the local maxima with a value larger than zero. If more than one local peak were detected for a signal segment, the local peak was defined as the one with the largest peak amplitude. We focused on the delay profiles with at least 102 (128 × 80%) local peaks within a time segment. For those selected delay profiles, if the local peak was less than zero or no local maxima was detected, its relative time delay with respect to the global peak was defined as the mean of its three neighboring electrodes. Next, a delay matrix was formed by concatenating all of the delay profiles, to which we then applied SVD to extract the principal delay profile.

The ECoG gamma-power signals were projected onto the principal delay profile direction to identify time segments with propagations. The time segments were generated as described above and only those with a global involvement, which account for 44.05 ± 3.59 % (mean ± SD across four monkeys) of the total segments, were analyzed. For each time segment, the Pearson’s correlation between the relative timing and position of local peaks across all the electrodes was calculated. To create a threshold for detecting time segments with propagations, we generated a null distribution of time-position correlations along the randomly rotated principal delay profiles with retaining the relative positions of electrodes which were obtained by randomly rotating the principal delay profiles on a coordinate plane with random degrees with respect to the x, y axes. The time segments with a bottom-up (from the SM regions to the high-order regions) propagation along the principal delay profile were identified if the time-position correlation had a positive value exceeding the threshold (1.64SD of the null distribution of time-position correlations). The time segments with a top-down (from the high-order regions to the SM regions) propagation along the principal delay profile were identified if the time-position correlation had a negative value exceeding the threshold (1.64SD of the null distribution of time-position correlations). The averaged top-down and bottom-up propagation maps were calculated as follows. First, we identified the global peaks for the time segments with propagations. Then, a 12.2-sec time window (61 × 0.2 sec; the temporal resolution of bandlimited power signals is 0.2 seconds) centering on each of these global peaks was defined to cover thirty time points before and thirty time points after the global peak. Second, we averaged the ECoG bandlimited power signals and associated time-position graphs of these time windows to obtain the averaged propagating maps.

Considering that SVD components captured the major propagating directions in an arbitrary unit, we rescaled the derived principal delay profile to second unit using the same strategy as we used for human rsfMRI data. Briefly, we calculated the regression coefficient of the position-time relationship for each segment with a propagation to estimate the propagating speed. Then the averaged propagating speed across all the time segments with propagations was computed and utilized to re-scale the principal delay profiles into the unit of seconds.

We quantified the cross-hierarchy pattern of the principal delay profile from monkeys by comparing it with the cortical myelination map, which has been suggested to be a good estimation of cortical anatomical hierarchy (36). Given that the cortical myelination map was available on the average Yerkes19 macaque surface (35), we manually mapped the location of 128 electrodes of each monkey onto the average Yerkes19 macaque surface (35) based on the gyrus and sulci of the brain (see **Fig. S8**). Next, we extracted a vector of the myelination values at the locations of the 128 electrodes for each monkey. Then, a Pearson’s correlation between the principal delay profile and this myelination vector was calculated to estimate the spatial similarity of the two, which was used for quantifying the cross-hierarchy pattern of the principal delay profile.

The principal delay profile was also calculated from ECoG gamma powers with skipping the temporal filtering and/or spatial filtering, or switching their order (**Fig. S15**).

### Fine-scale propagations within sensory modalities in the rsfMRI signals

A simple linear regression was applied to examine the relationship between the delay in the principal delay profile and the hierarchy level across brain regions within each sensory modality. Each vertex was numbered according to the hierarchy level of the brain region to which the vertex belongs. In the simple linear regression model, the hierarchy level of vertices was the predictor variable and the delay value was the response variable. A significant p-value of the regression coefficient would indicate whether the propagation is along the hierarchy order in a sensory modality.

The hierarchical level of different visual regions was determined according to (39). The hierarchy of the auditory system was determined according to (81, 82). The hierarchy of the somatosensory cortex was determined according to (83). The retinotopy map (84) was used to identify the peripheral and foveal areas of the V1-V4. Peripheral and foveal areas in the V1-V4 were divided based on a cut-off value of 2.5 in the eccentricity map. The atlas of topographic subareas in the somatosensory-motor strip (85) was used to identify brain regions responsible for face, upper limb, trunk, lower limb, and eye. The HCP’s multi-modal cortical parcellation atlas was used to locate different brain regions on the cortical surface. All these atlases can be downloaded from the website (https://balsa.wustl.edu).

### Subcortical co-activations/de-activations associated with rsfMRI propagations

The temporal dynamics of subcortical regions at the top-down and bottom-up propagations were studied mainly based on the z-score maps of the propagations in the volume space, which were obtained as described above. We simply averaged these z-score maps at various time points across voxels within any subcortical region of interests. The location of thalamic nuclei/regions was determined according to Morel Atlas (86). The Harvard ascending arousal network (AAN) atlas was used to locate the brainstem nuclei of the ascending arousal network (87). The masks of the substantia nigra (SN), the nucleus accumbens (NAc), and the nucleus basalis (NB) were acquired by taking the overlap between the brain regions showing significant (Z < -7) de-activations at time zero z-score map for the bottom-up propagation and those defined by brain atlases of SN (88), NAc (80) and NB (6). The de-activations in SN and NAc appeared to be only in a subsection of atlas-defined structures (**Fig. S20**).

### Modulation of cross-hierarchy propagations across different brain states

We quantified and compared the occurrence rate of the top-down and bottom-up propagations across different brain states or sessions that are likely associated with distinct arousal levels. For the human rsfMRI data, we classified fMRI sessions into sub-groups with different arousal levels, which were estimated in two different ways. First, we adapted a template-matching method (42) to estimate the arousal level based on a previous study (89). Briefly, we calculated the spatial correlation between individual rsfMRI time points and a global co-activation pattern that has been linked to transient arousal events previously (6). Then, we extracted the envelope amplitude of this spatial correlation time course and averaged it within each session to quantify the general arousal state of subjects in a specific session. The envelope amplitude was computed as the absolute value of the Hilbert transform. Since the presence of this global co-activation pattern suggest the occurrence of transient arousal events and thus relatively drowsy state, a higher value of this fMRI-based metric is corresponding to a lower arousal state. Based on this fMRI-based arousal measure, we divided all of the rsfMRI scanning sessions into three groups of equal size (*N* = 613 sessions for each, each session is 15 minutes in length) with low, medium, and high arousal levels, and then compared the occurrence rate of the rsfMRI propagations. Secondly, we identified, based on the note of HCP experimenters, a subset of 117 rsfMRI sessions in which the subjects were noted to be sleeping (90). We then compared the occurrence rate of propagations in this subset and all other sessions (N = 1723 sessions). For the ECoG gamma-band powers concatenated from the four monkeys, we compared the occurrence rate of the top-down and bottom-up propagations across sessions collected in three different conditions: 11 sessions (each session is 25 minutes in length) for eyes-open state, 10 sessions for eyes-closed state, 15 sessions for sleep state.

Since these propagations are detected only in rsfMRI/ECoG segments with a global involvement, and their occurrence rate changes across arousal states might simply reflect a change of the global signal, which has been shown to be closely related to brain arousal level (91–95). For this reason, we also calculated and compared the ratio of the top-down propagations to the bottom-up propagations that should be exempted from any potential bias caused by the change of globally averaged signal.

The statistical significance of the difference in the occurrence rate of propagations, as well as its ratio, across different arousal states were determined using permutation test. For each comparison described above, we randomly divided the data into subgroups of the same size, calculated the differences, and repeated the procedure 1,000,000 times to build a null distribution for the differences. Then, we obtained the p-values of the observed differences by comparing them with the null distributions.

## Acknowledgments

We thank Dr. Matthew F. Glasser for sharing the information about subjects’ sleeping state during the scanning and the Computer Vision Laboratory for sharing the Morel thalamus atlas. We also thank Dr. Yunjie Tong for sharing the averaged time lag map from dynamic susceptibility contrast. This research was supported by the NIH Pathway to Independence Award (K99/R00) 5R00NS092996-03. The human data were analyzed using the computing resources provided by the Institute for Computational and Data Sciences at the Pennsylvania State University (https://icds.psu.edu). The human data were provided by the Human Connectome Project, WU-Minn Consortium (Principal Investigators: David Van Essen and Kamil Ugurbil; 1U54MH091657) funded by the 16 NIH Institutes and Centers that support the NIH Blueprint for Neuroscience Research; and by the McDonnell Center for Systems Neuroscience at Washington University.

## Author contributions

Y.G. and X.L. designed the study and performed the analyses; Y.G., L.S., S.K., and J.W. draw the figures; S.K. and Y.G. mapped ECoG electrodes onto the average macaque surface; all authors contributed to editing the paper.

## Competing interests

Authors declare no competing interests.

## Data and materials availability

Both the HCP data and the monkey electrophysiology dataset are public and available from the websites https://www.humanconnectome.org and http://neurotycho.org/anesthesia-and-sleep-task respectively. The MATLAB code for the analysis and the human principal delay profile in this study are available at GitHub: https://github.com/YamengGu/the-cross-hierarchy-propagation (96).

## List of Supplementary Materials

Figures S1-S20.

## Legends for Supplementary Animations

**Animation 1:**
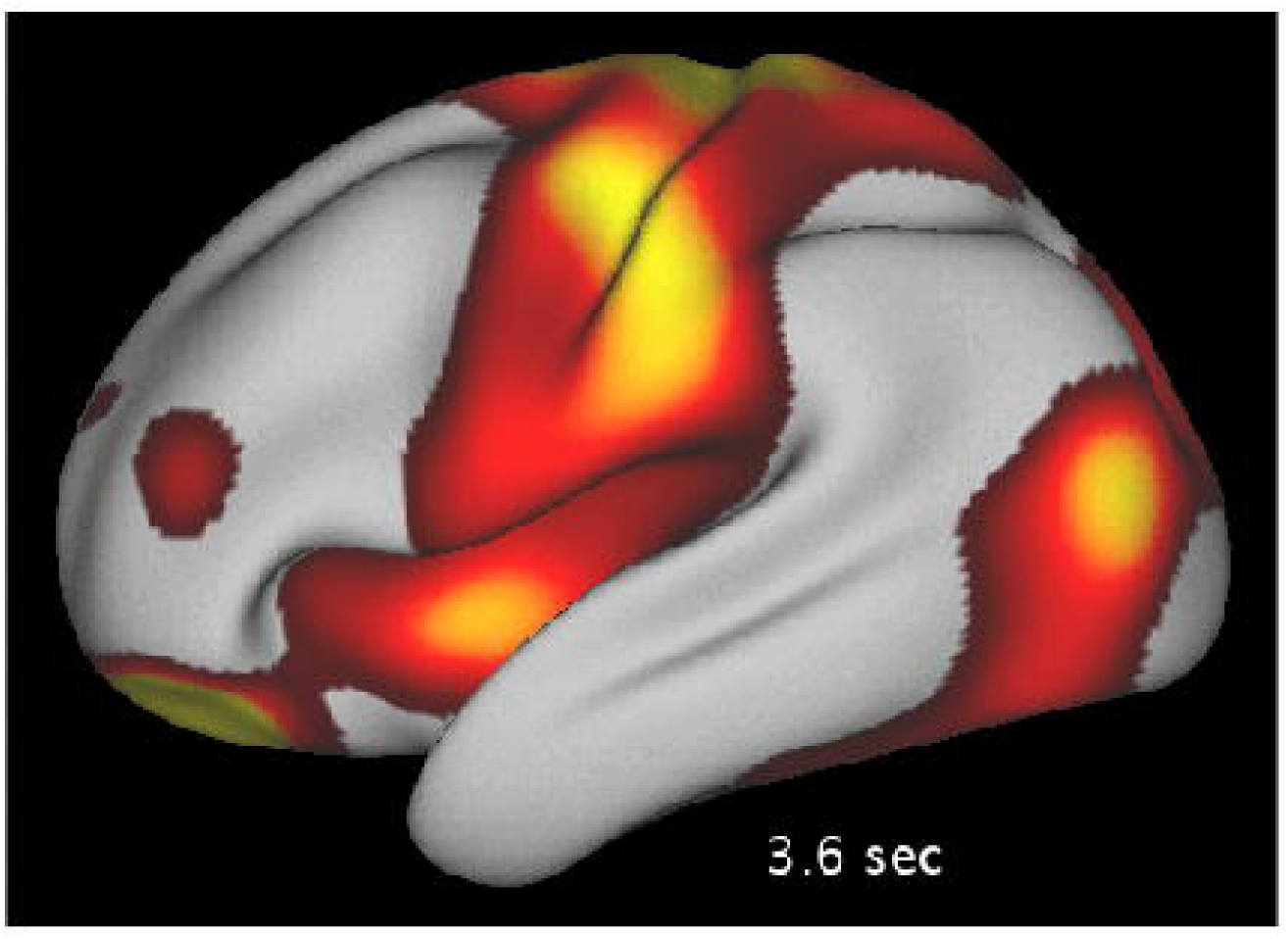
Single exemplary instance of bottom-up propagation in human rsfMRI on the brain surface.

**Animation 2:**
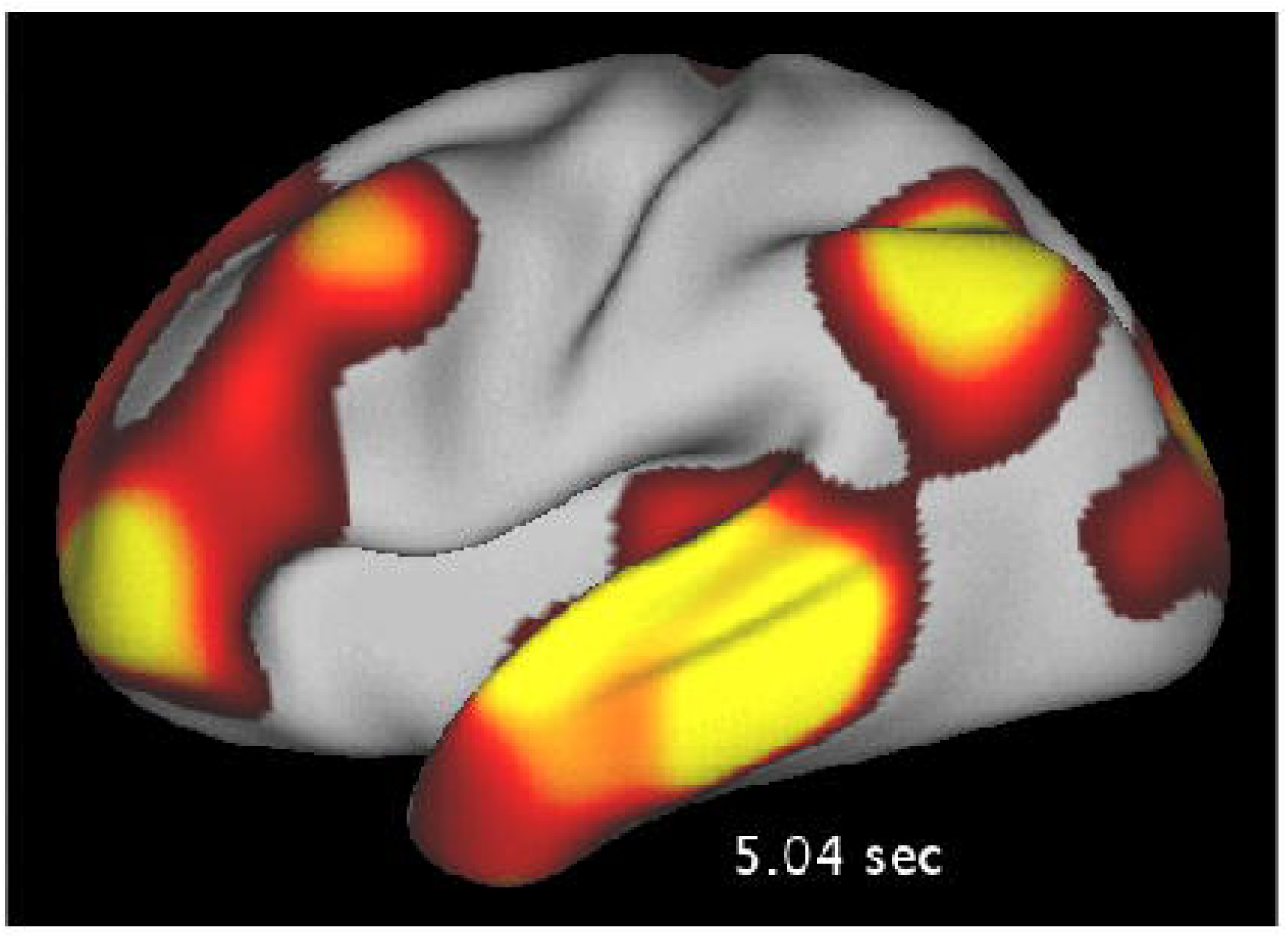
Single exemplary instance of top-down propagation in human rsfMRI on the brain surface.

**Animation 3:**
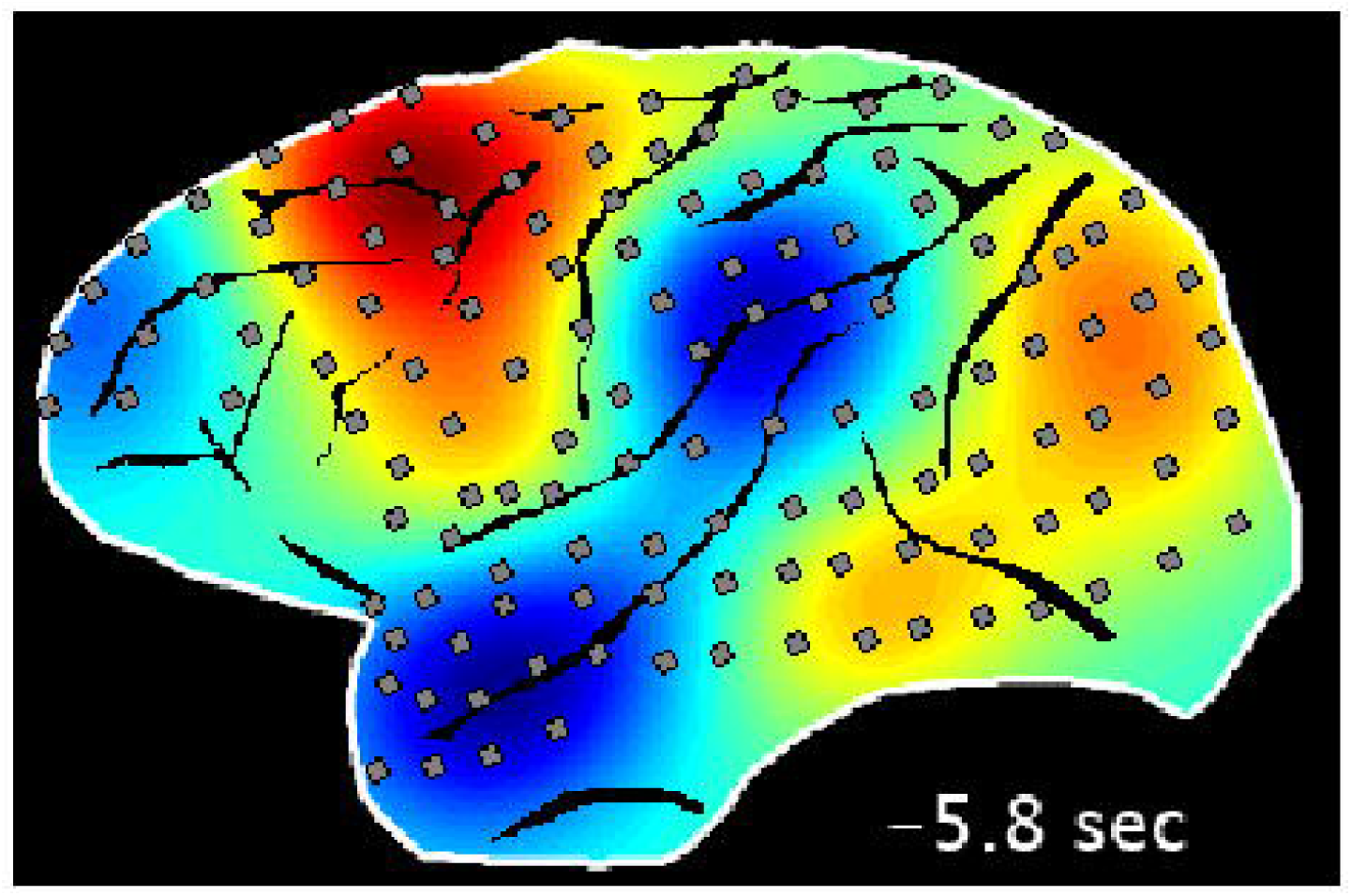
Single exemplary instance of bottom-up propagation in monkey ECoG on the brain surface.

**Animation 4:**
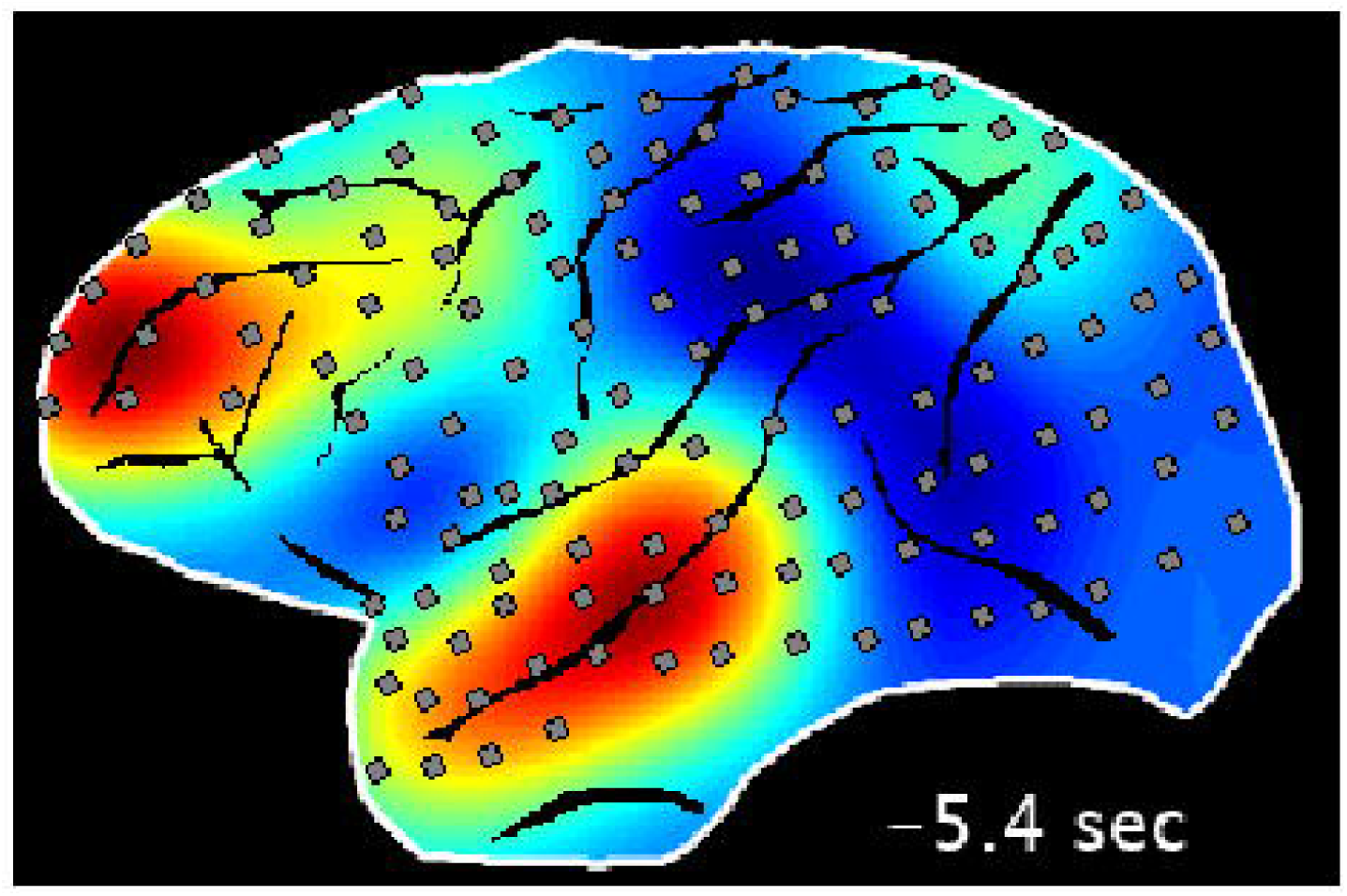
Single exemplary instance of top-down propagation in monkey ECoG on the brain surface.

## Notes

### Competing Interest Statement

The authors have declared no competing interest.

