## Supplementary material for "Brain activity fluctuations propagate as waves traversing the cortical hierarchy": Fig. S1, Fig. S2, Fig. S3, Fig. S4,Fig. S5, Fig. S6, Fig. S7, Fig. S8, Fig. S9, Fig. S10, Fig. S11, Fig. S12, Fig. S13, Fig. S14, Fig. S15, Fig. S16

**This file includes:**

Figs. S1 to S20.

**Figures S1-S20**

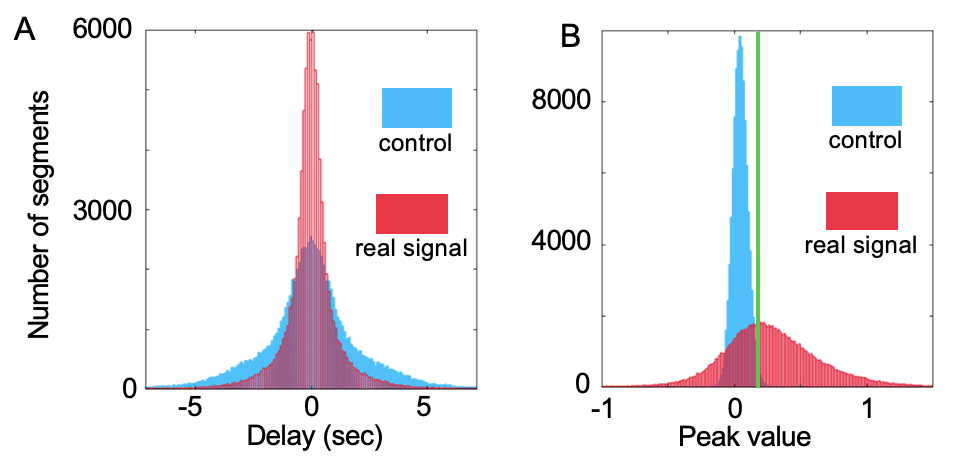

Fig. S1.

Detecting rsfMRI segments with a global involvement. The active band observed in Fig. 1B indicated that the local peaks of different time points within a time segment are clustered together and the phase of local peaks are synchronized. To disrupt the synchronization, we randomly shifted the time series of rsfMRI signal and the global peak value within time segments were recalculated to build a null model as the control group shown in blue. As expected, the delay between the local peak and the global mean peak within segments in real signal are smaller compared to the null distribution (A) and the global peak values are larger in real signal compared to the near-zero values in control group (B). The distribution of delay and peak value from real signal and control group are significantly different (A: *p* = 0, two-sample Kolmogorov-Smirnov test; B: *p* = 0, two-sample Kolmogorov-Smirnov test). The segments with a global involvement (58.80%) are detected if their global signal peak values exceed 99^th^ percentile of null distribution (green vertical line in B).

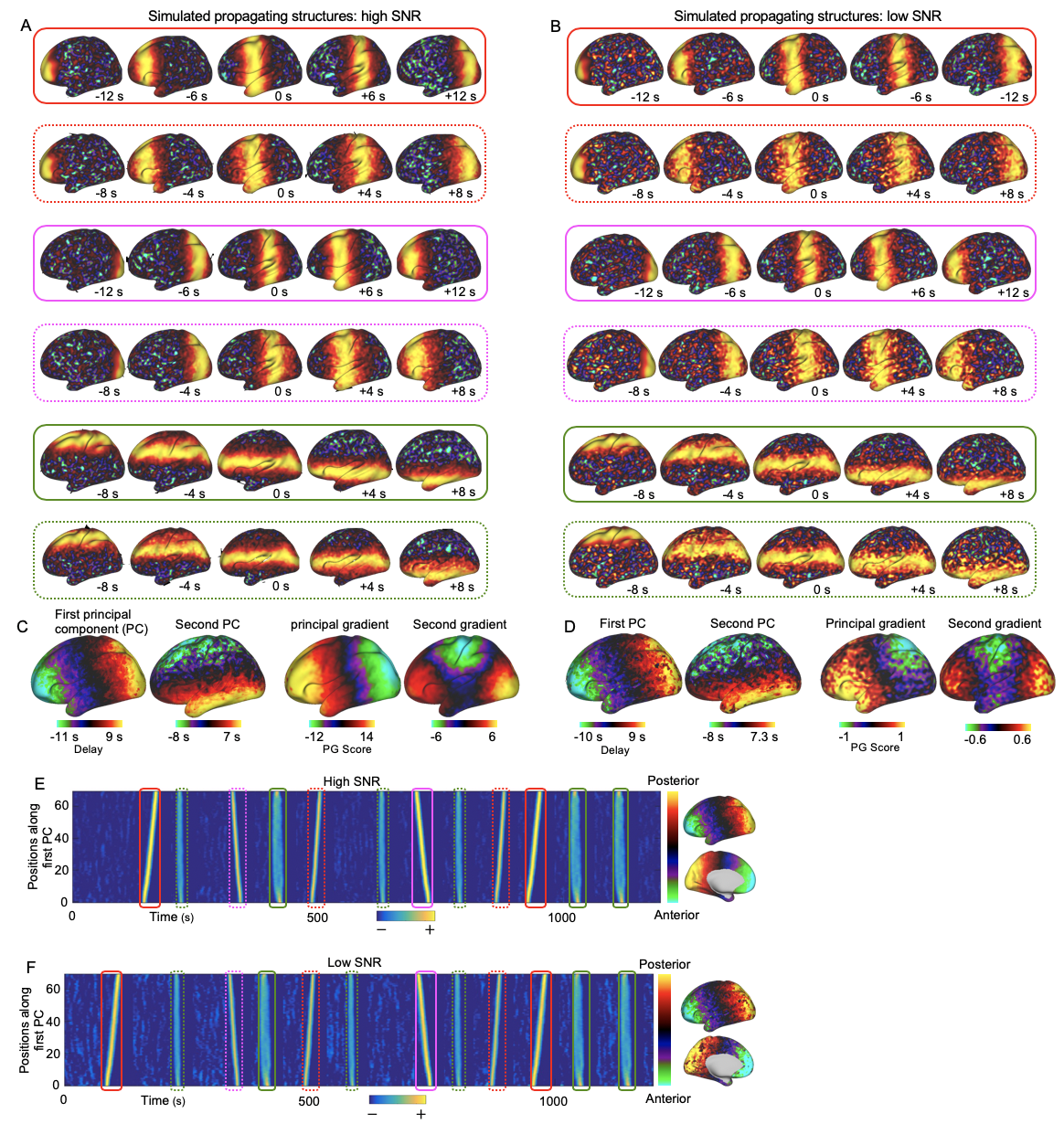
**Fig. S2.**

The method of decomposing delay profiles can better recover propagating directions than the PG method of embedding the rsfMRI connectivity matrix into a low-dimensional space. We simulated two types of propagating structures along the anterior-posterior direction, posterior-anterior direction, and dorsal-ventral direction, one having a high signal-to-noise ratio (SNR) (A), i.e., the level of the active band to the level of background noise, and the other one having low SNR (B). The time zero is when the active band was located at the center of brain surface. Then the simulated propagating structures were randomly inserted into simulated rsfMRI signal modeled as white noise, generating time series with propagating structures. Applying the method of decomposing delay profiles to both high SNR (2SD, A) and low SNR (5SD, B) simulated time series successfully recovered two propagating directions in (C, left) and (D, left) respectively, indicated by the first two principal components. In contrast, the PG method only recovered one direction in the high SNR simulated data (C, right) and failed to recover any directions in the low SNR simulated data (D, right). Projecting an exemplary time series along the principal delay profile (the first principal component) generated the time-position graphs and showed clear tilted bands in both high SNR (E) and low SNR simulated data (F), indicating the existence of propagating structures along the anterior-to-posterior axis.

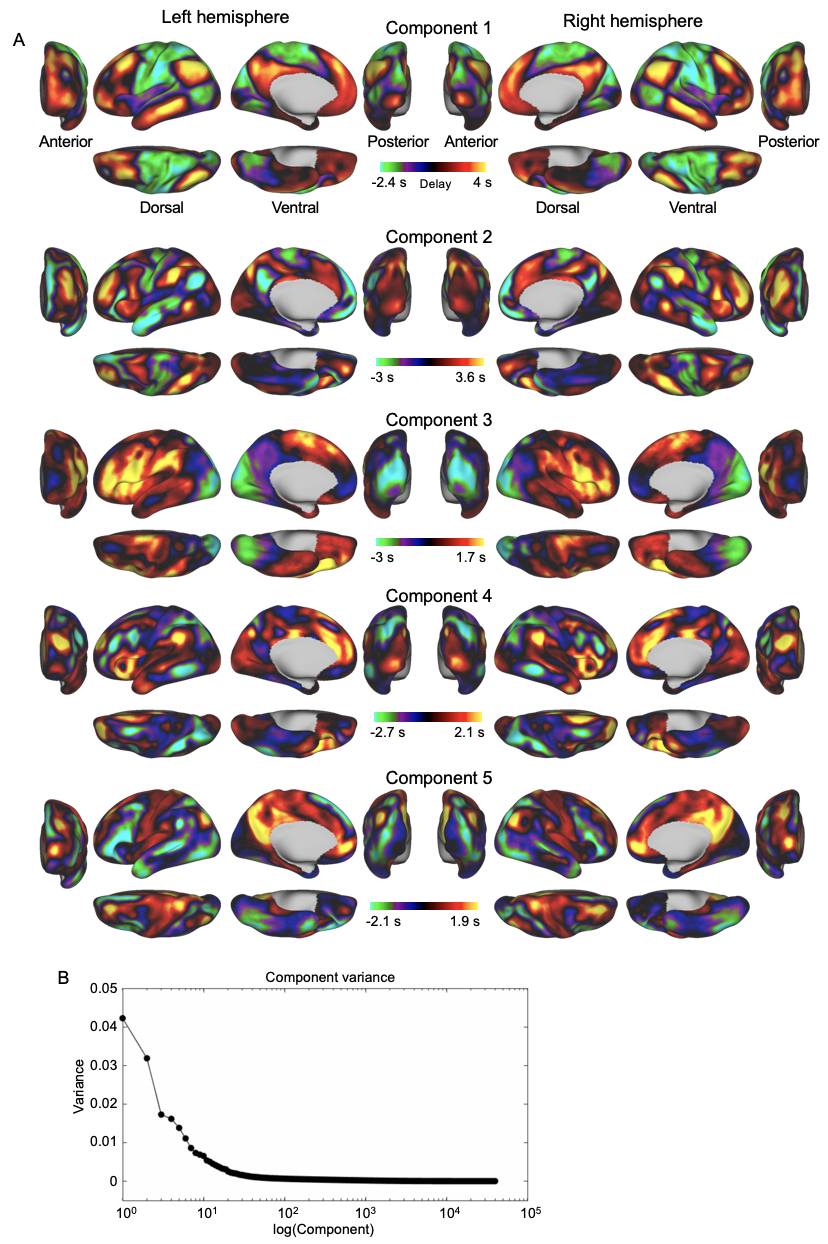

**Fig. S3.**

The first 5 principal components (A) explaining the largest variance (B) extracted by applying the method of decomposing delay profiles on all of the four rsfMRI sessions of 460 human subjects.

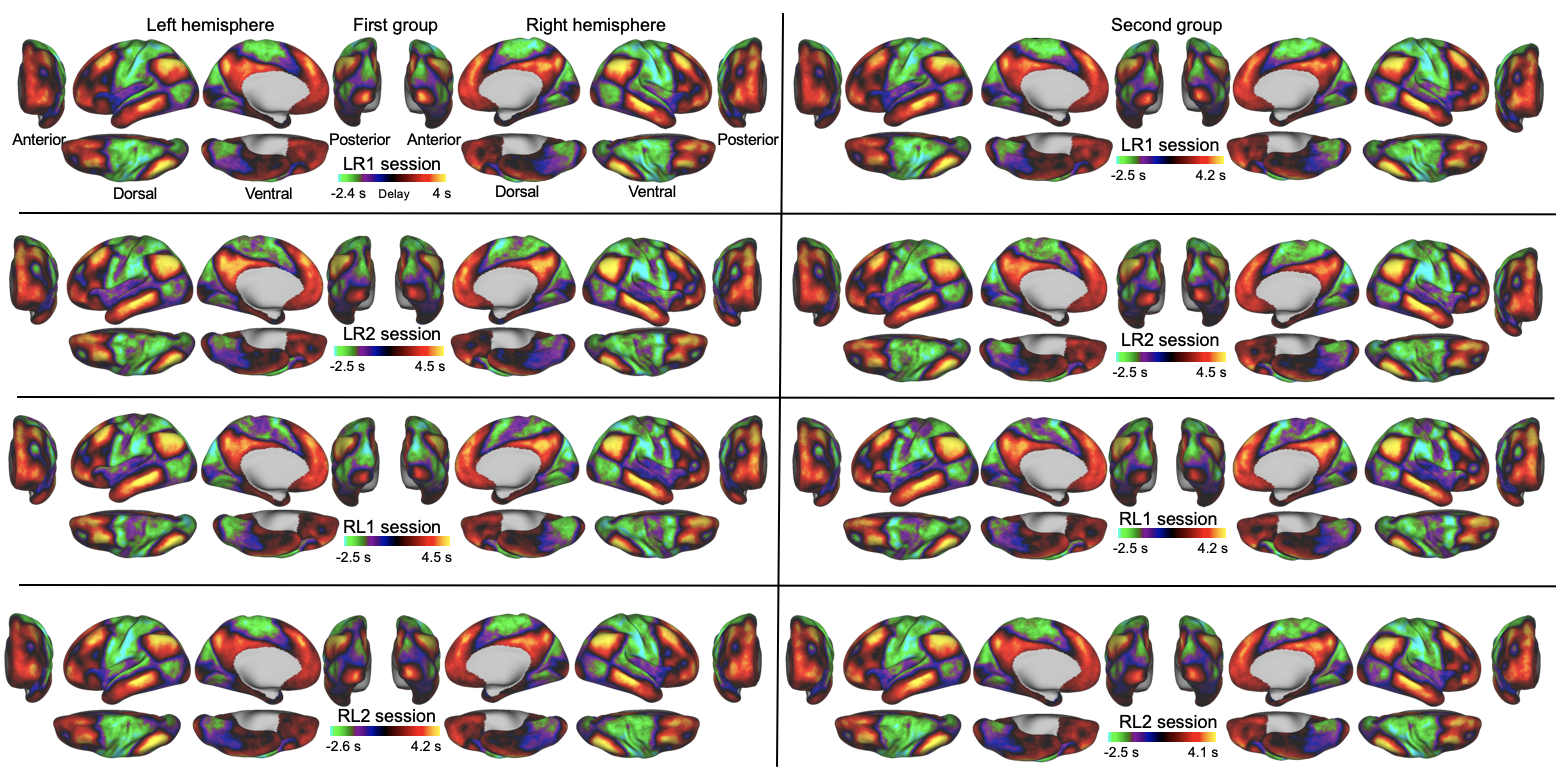

**Fig. S4.**

The reproducibility of the principal propagating direction along the cross-hierarchy in human rsfMRI. The split-half analyses were applied on each of the four scanning sessions (LR1, LR2, RL1, RL2). Specifically, we randomly split subjects in each scanning session into two equal groups and then applied the method of decomposing delay profiles on each group to extract the first principal component, i.e., the principal delay profile representing the principal propagating direction. All of the computed principal delay profiles showed a cross-hierarchy contrast, similar to the principal delay profile computed from all the four sessions in Fig. 2C. Left: the principal delay profile of the first group from four sessions. Right: the principal delay profile of the second group from four sessions. Each row represents the principal delay profile of the two groups in each session.

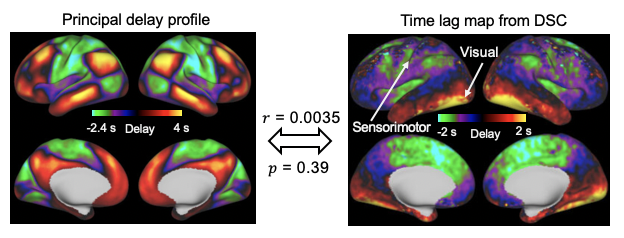

**Fig. S5.**

A comparison between the principal delay profile of human rsfMRI and the vascular lag map. Left: the principal delay profile from human rsfMRI. Right: the averaged time lag map derived from the dynamic susceptibility contrast (DSC) MRI scan (1). The spatial correlation between the two maps is 0.0035 (*p* = 0.39).

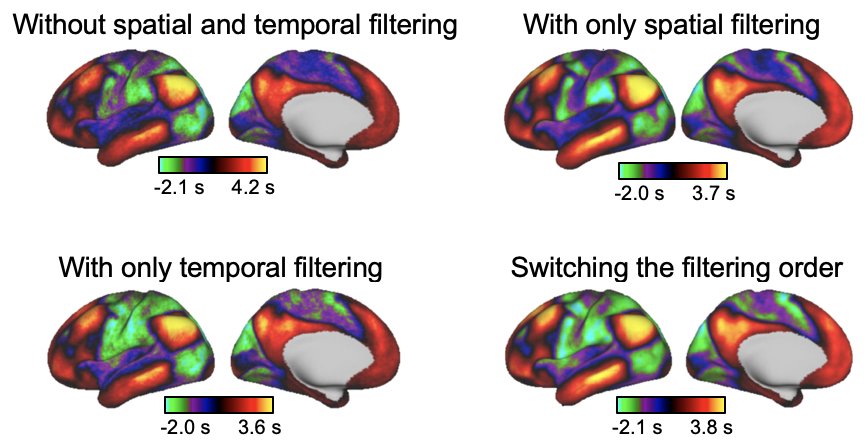

**Fig. S6.**

The effects of spatial and temporal filtering on the human fMRI. The principal delay profiles were computed without spatial and temporal filtering, with either spatial or temporal filtering, and with switching the order of filtering, i.e., first applying the temporal filtering and then the spatial filtering, which is opposite to that in Fig. 2.

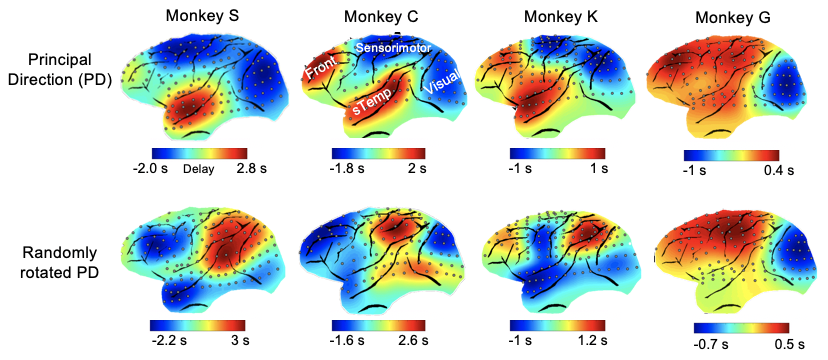

**Fig. S7.**

The reproducibility of the principal propagating direction along the cross-hierarchy in ECoG gamma-band power. Top panel: the principal delay profile of the ECoG gamma-band (42–95 Hz) power from four monkeys. Bottom panel: the corresponding randomly rotated principal delay profile from four monkeys. Note that all of the principal delay profiles are the first principal component generated by applying the method of decomposing delay profiles except that the one in Monkey S is the second principal component.

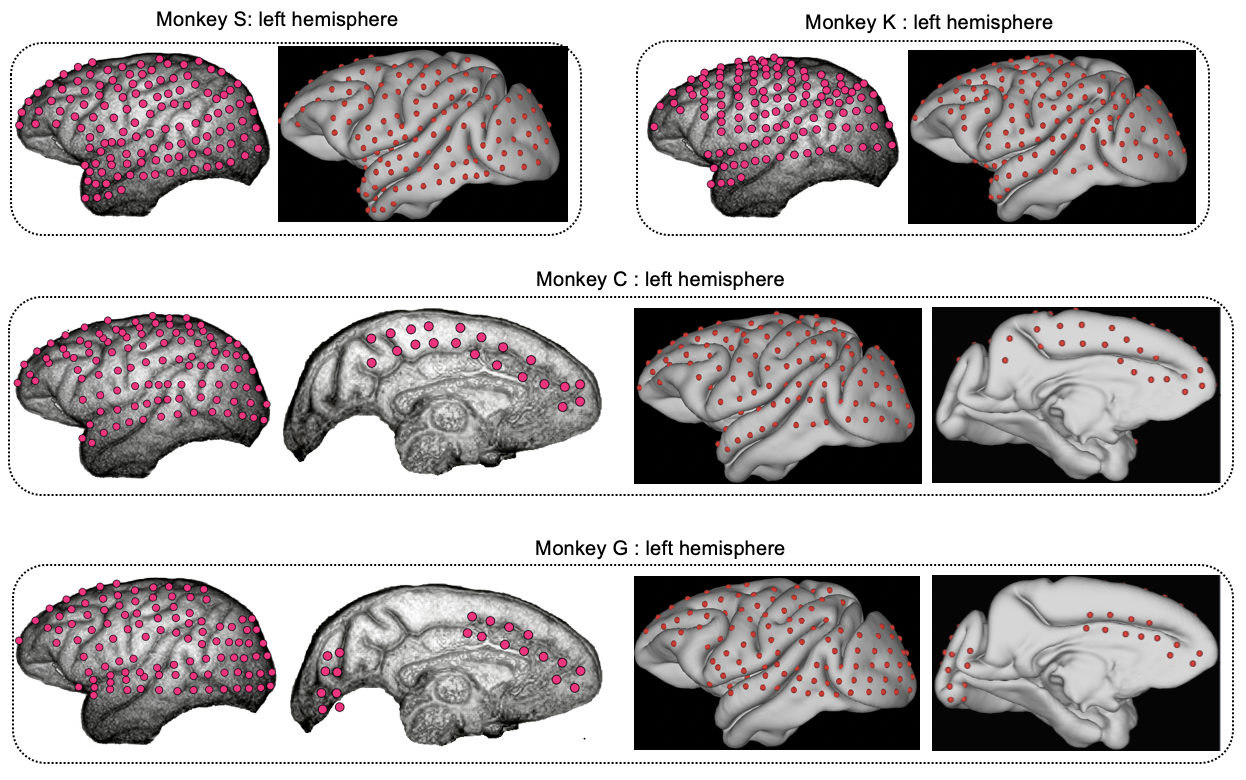

**Fig. S8.**

The raw 128 electrodes (magenta) of four monkeys were manually mapped onto the average Yerkes19 macaque surface (red electrodes) based on the gyrus and sulci of the brain.

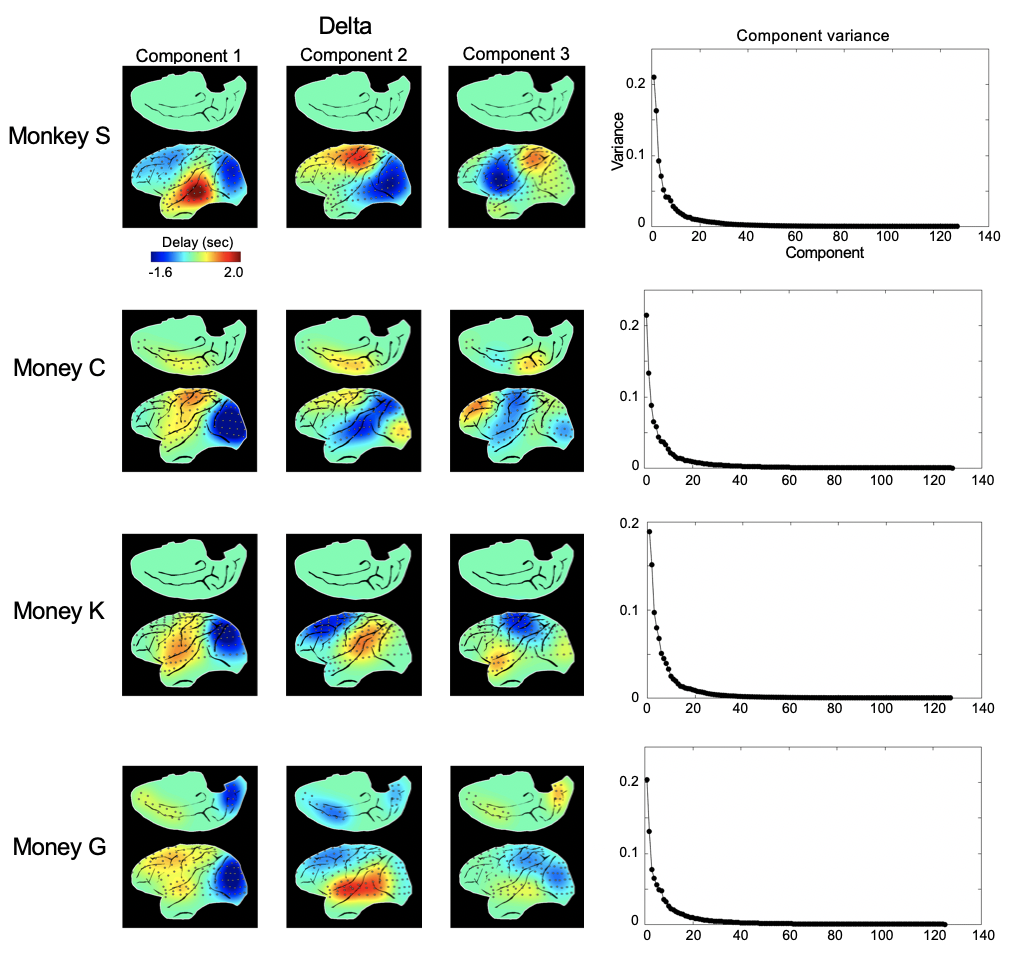

**Fig. S9.**

The first three principal components explaining the largest variance extracted by applying the method of decomposing delay profiles within the delta frequency band (1–4 Hz) for each monkey. Right: the variance explained by each component.

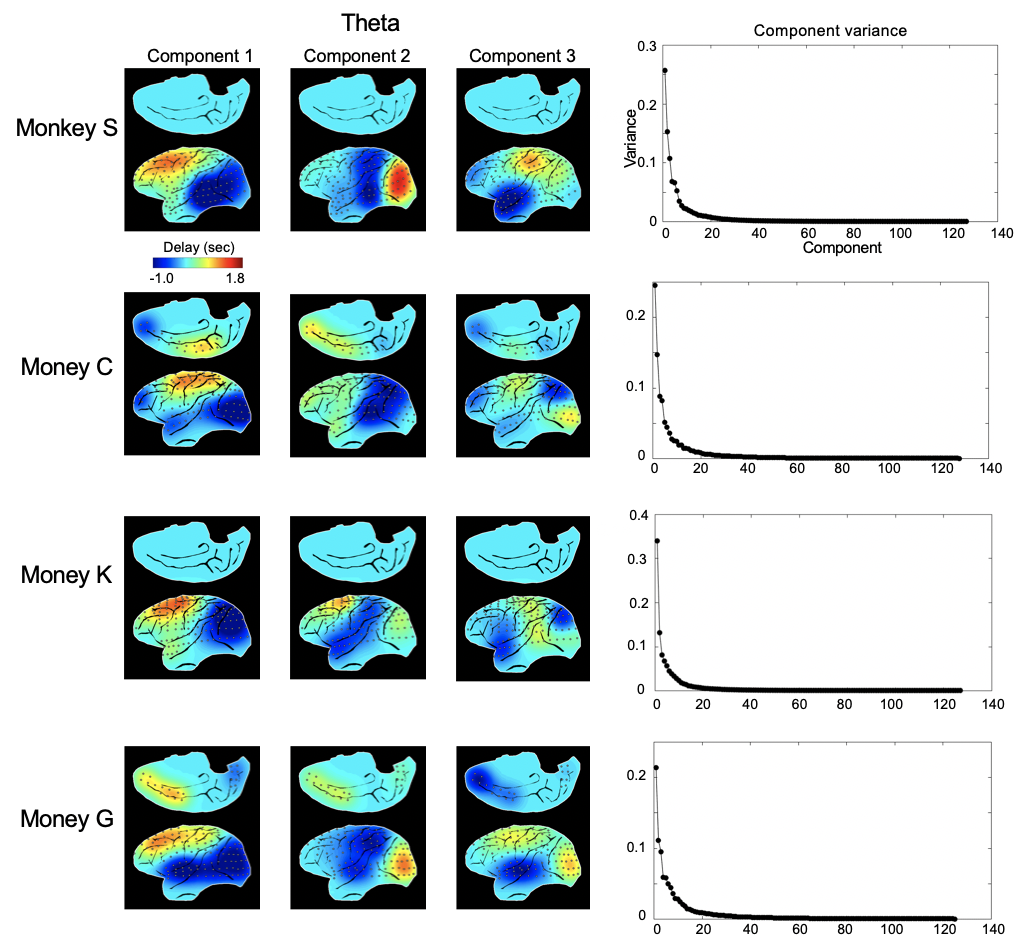

**Fig. S10.**

The first three principal components explaining the largest variance extracted by applying the method of decomposing delay profiles within the theta frequency band (5–8 Hz) for each monkey. Right: the variance explained by each component.

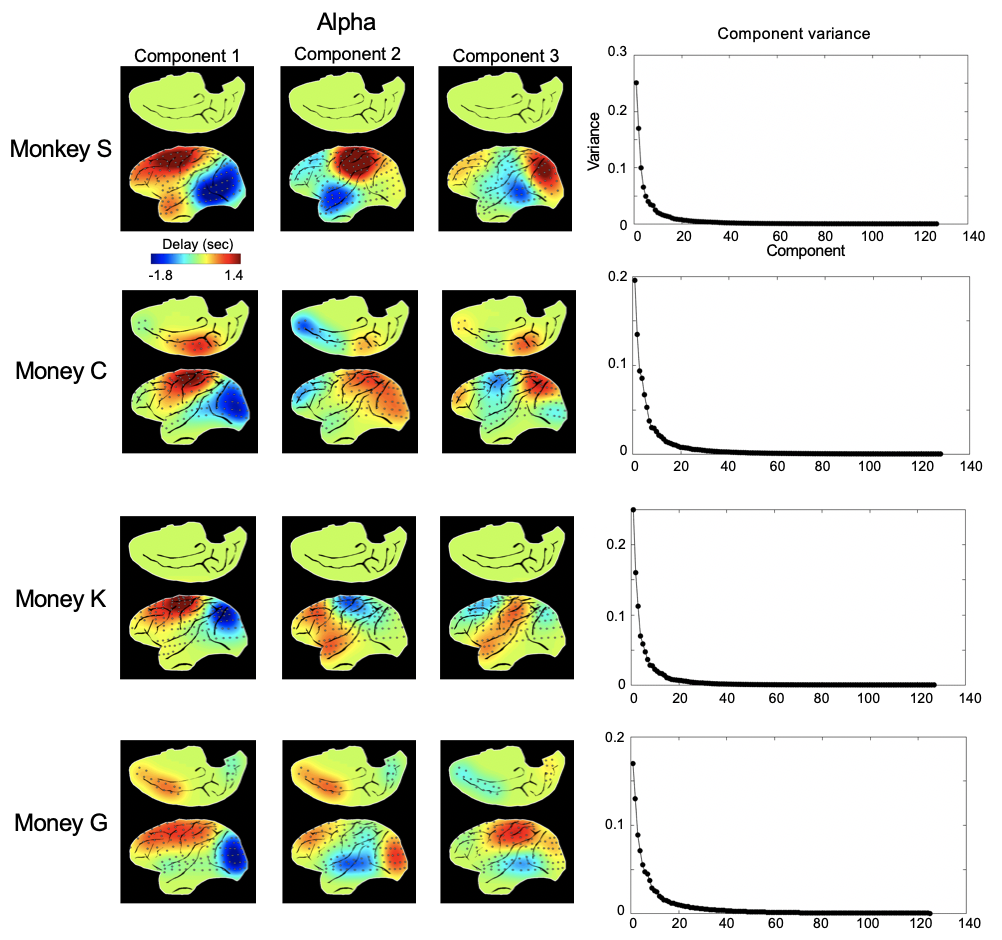

**Fig. S11.**

The first three principal components explaining the largest variance extracted by applying the method of decomposing delay profiles within the alpha frequency band (9–15 Hz) for each monkey. Right: the variance explained by each component.

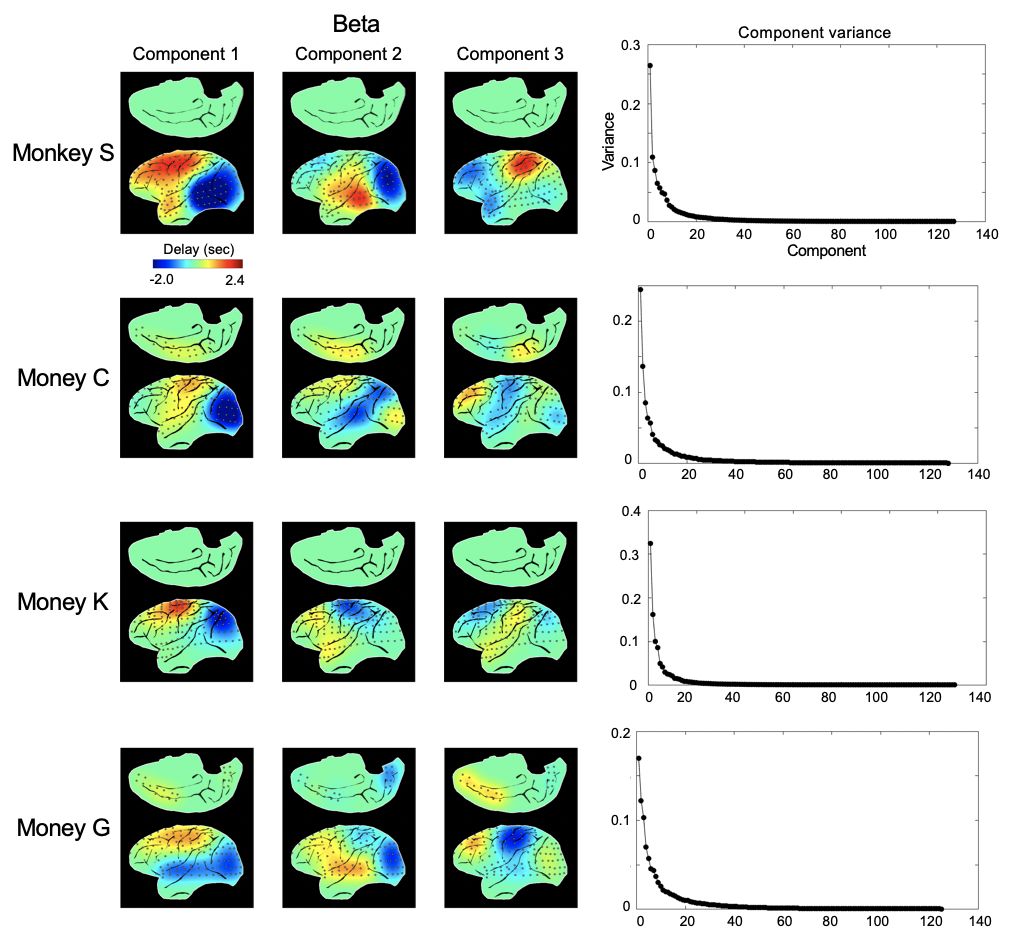

**Fig. S12.**

The first three principal components explaining the largest variance extracted by applying the method of decomposing delay profiles within the beta frequency band (17–32 Hz) for each monkey. Right: the variance explained by each component.

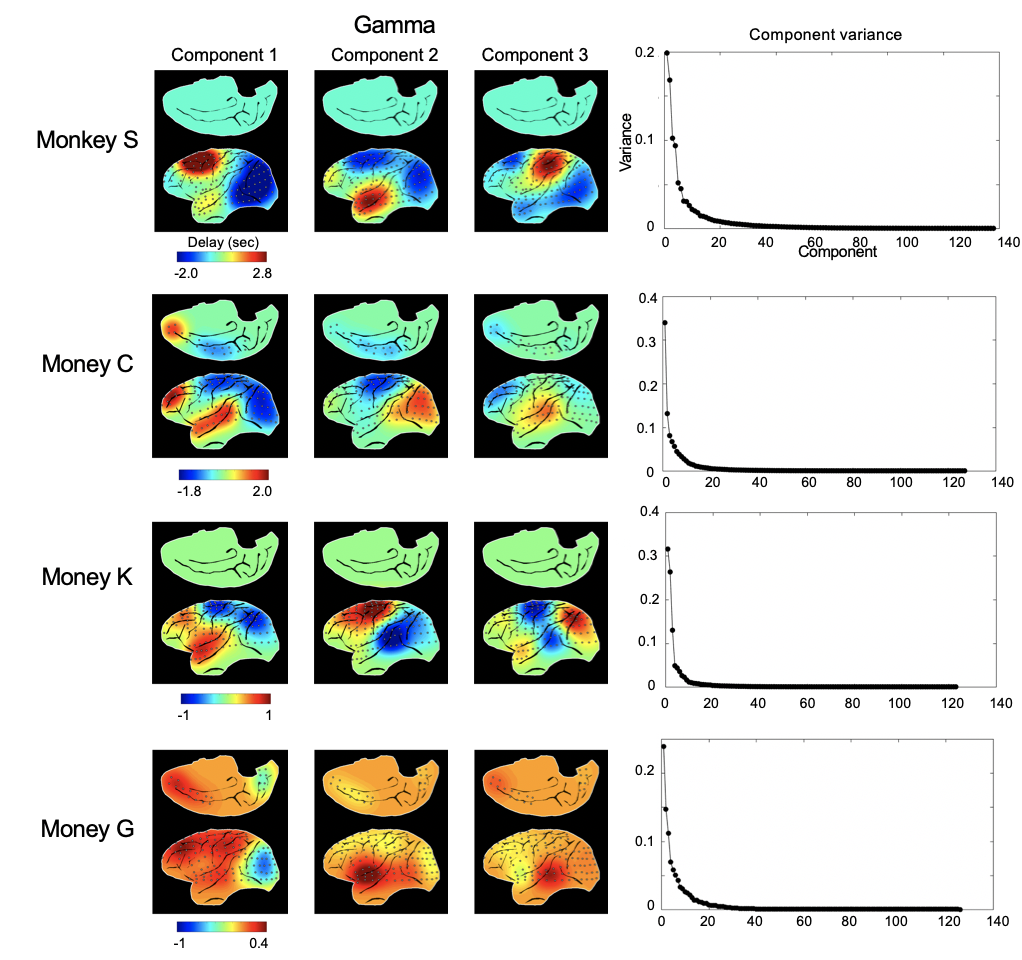

**Fig. S13.**

The first three principal components explaining the largest variance extracted by applying the method of decomposing delay profiles within the gamma frequency band (42–95 Hz) for each monkey. Right: the variance explained by each component.

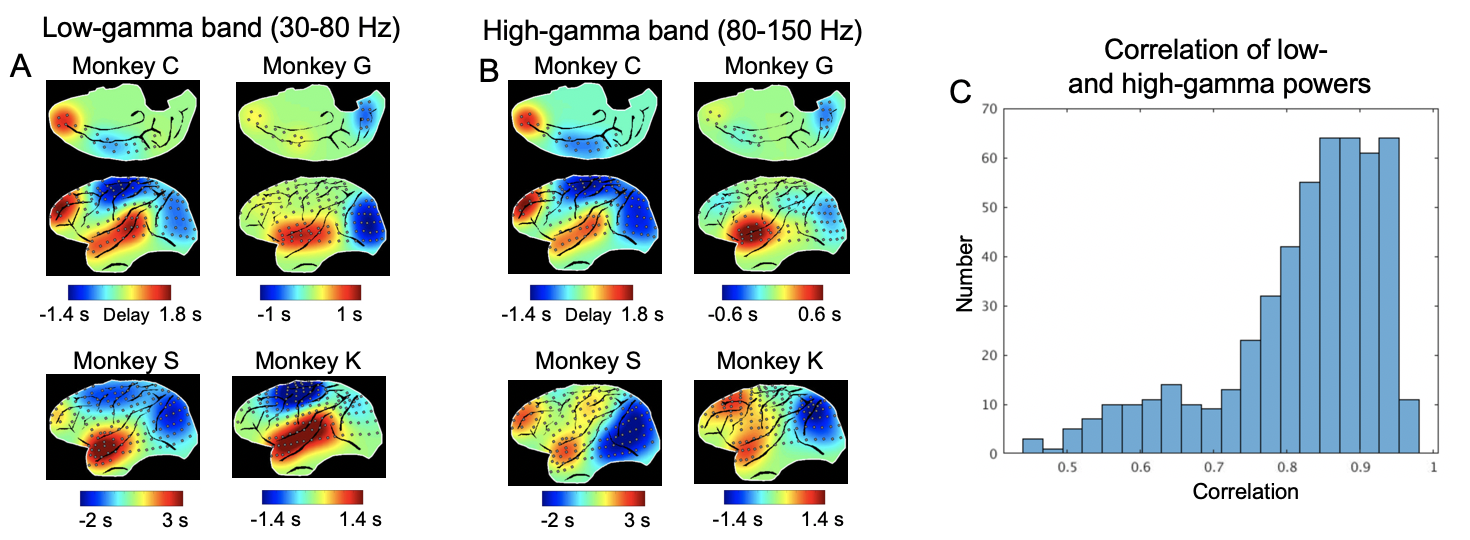

**Fig. S14.**

The principal delay profiles of two sub-bands of the gamma range. (A) The principal delay profile of the low-gamma band (30-80 Hz) power in monkey ECoG shows a cross-hierarchy contrast. (B) The similar result for the high-gamma band (80-150 Hz) power of monkey ECoG. (C) The temporal correlations between the low-gamma and high-gamma powers for all the electrodes from all 4 monkeys.

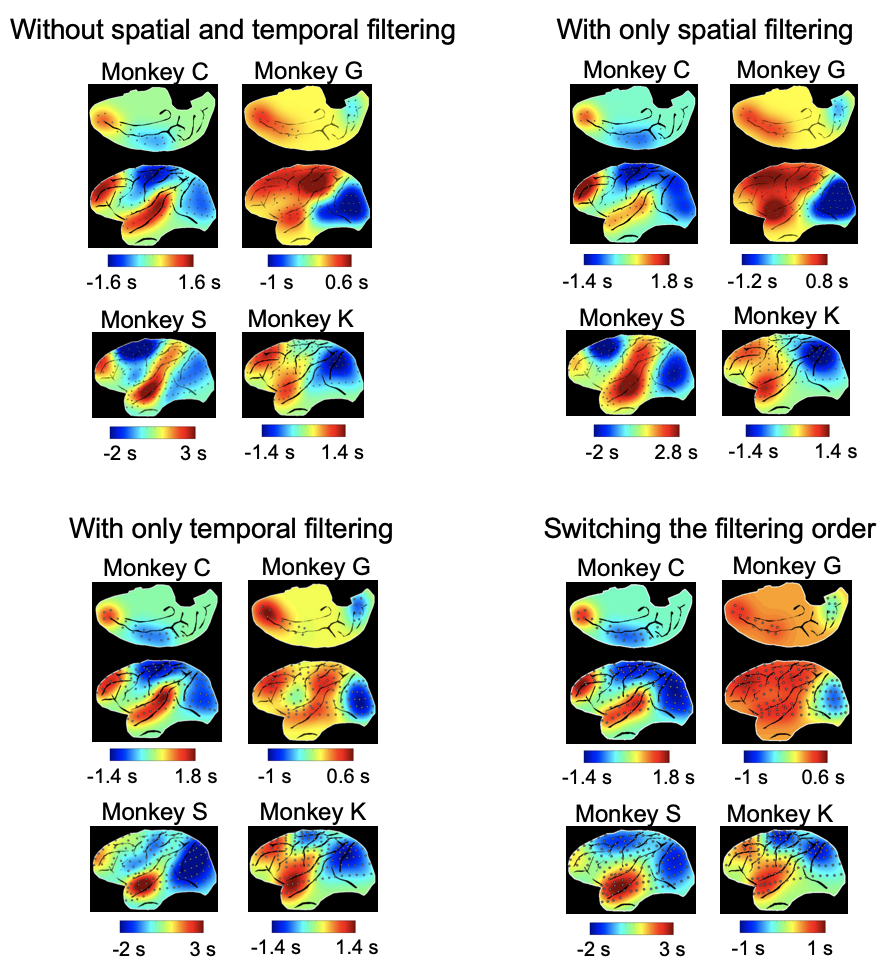

**Fig. S15.**

The effects of spatial and temporal filtering on the monkey ECoG results. The principal delay profiles were computed without spatial and temporal filtering, with either spatial or temporal filtering, and with switching the order of filtering, i.e., first applying the temporal filtering and then the spatial filtering, which is opposite to that in Fig. 3.

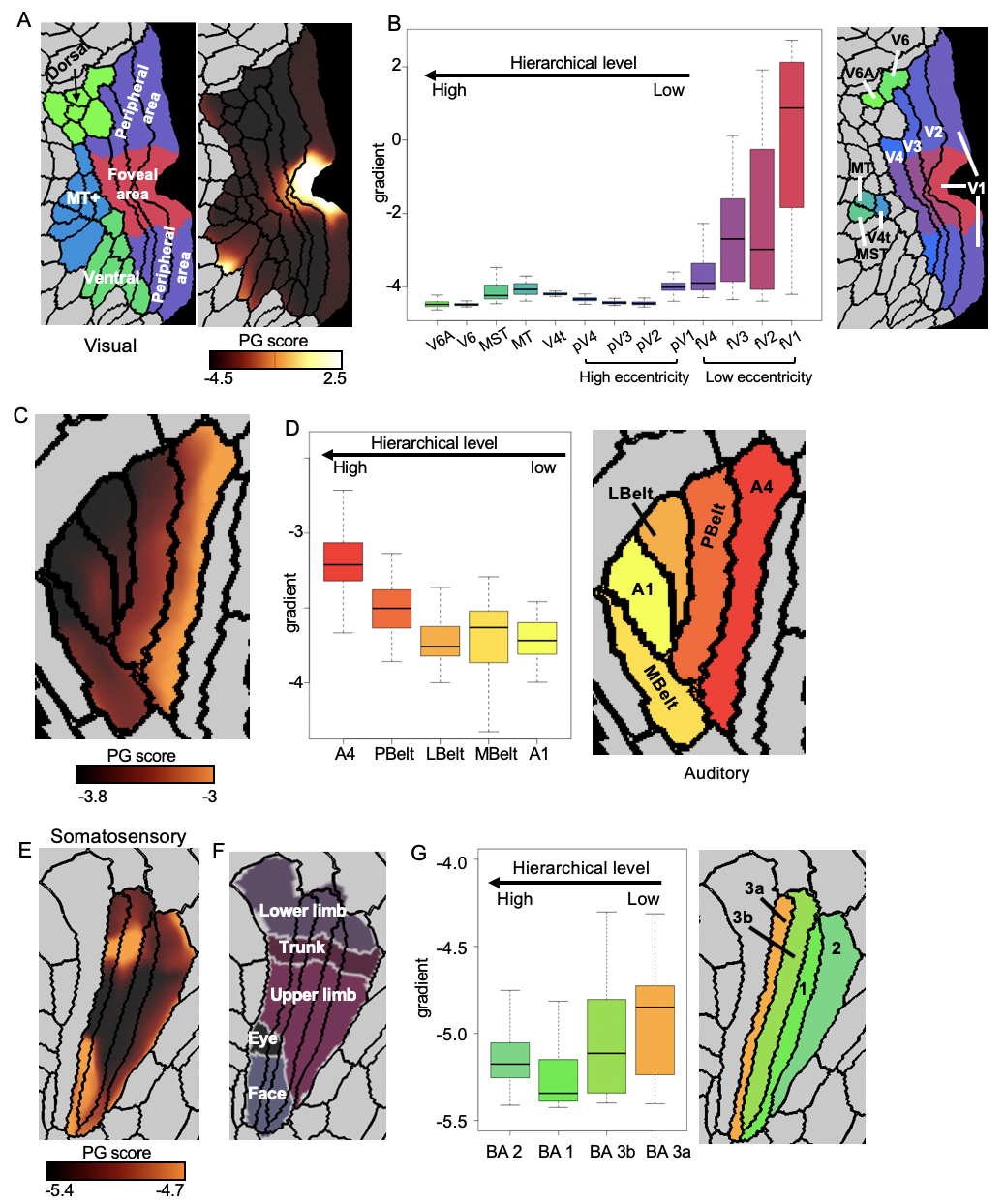

**Fig. S16.**

The relationship between the principal gradient (PG) score and the hierarchical level within each sensory modality. (A) The local contrast of the principal gradient within the three visual-related regions (MT+ complex, Dorsal stream, Ventral stream) defined by a multi-modal parcellation atlas (2). (B) The averaged PG score of 13 visual parcels that were arranged according to their hierarchical (3) and retinotopic relationships (4). The simple linear regression indicated a significant relationship between the PG score and the hierarchy level across brain regions (*p* = 5.5x10^-113^ for fV1-fV2-fV3-fV4-V4t-MT-MST-V6-V6A and *p* = 2.5x10^-253^ for pV1-pV2-pV3-pV4-V4t-MT-MST-V6-V6A). (C-D) Results for the auditory system indicated a similar contrast between the primary and association auditory areas. The simple linear regression indicated a significant relationship between the PG score and the hierarchy level across auditory regions (*p* = 2.8x10^-206^). (E-G) Results for the somatosensory system. (E-F) The contrast of the principal gradient also shows certain correspondence with the somatotopic arrangement (5). (G) The simple linear regression indicated a significant relationship between the PG score and the hierarchy level across somatosensory regions (*p* = 8.8x10^-66^). Abbreviation: Dorsal, dorsal stream. Ventral, ventral stream. V6A, area V6A. V6, sixth visual area. MT+, MT+ complex. MST, medial superior temporal area. MT/V5, middle temporal area/fifth visual area. V4t, V4 transition zone. pV4, peripheral fourth visual area. pV3, peripheral third visual area. pV2, peripheral second visual area. pV1, peripheral primary visual cortex. fV4, foveal fourth visual area. fV3, foveal third visual area. fV2, foveal second visual area. fV1, foveal primary visual cortex. V4, fourth visual area. V3, third visual area. V2, second visual area. V1, primary visual cortex. A4, auditory 4 complex. PBelt, parabelt complex. LBelt, lateral belt complex. MBelt, medial belt complex. A1, primary auditory cortex. BA2, Brodmann’s area 2. BA1, Brodmann’s area 1. BA 3b, Brodmann’s area 3b. BA 3a, Brodmann’s area 3b. FEF, frontal eye fields. IPS, intraparietal sulcus area.

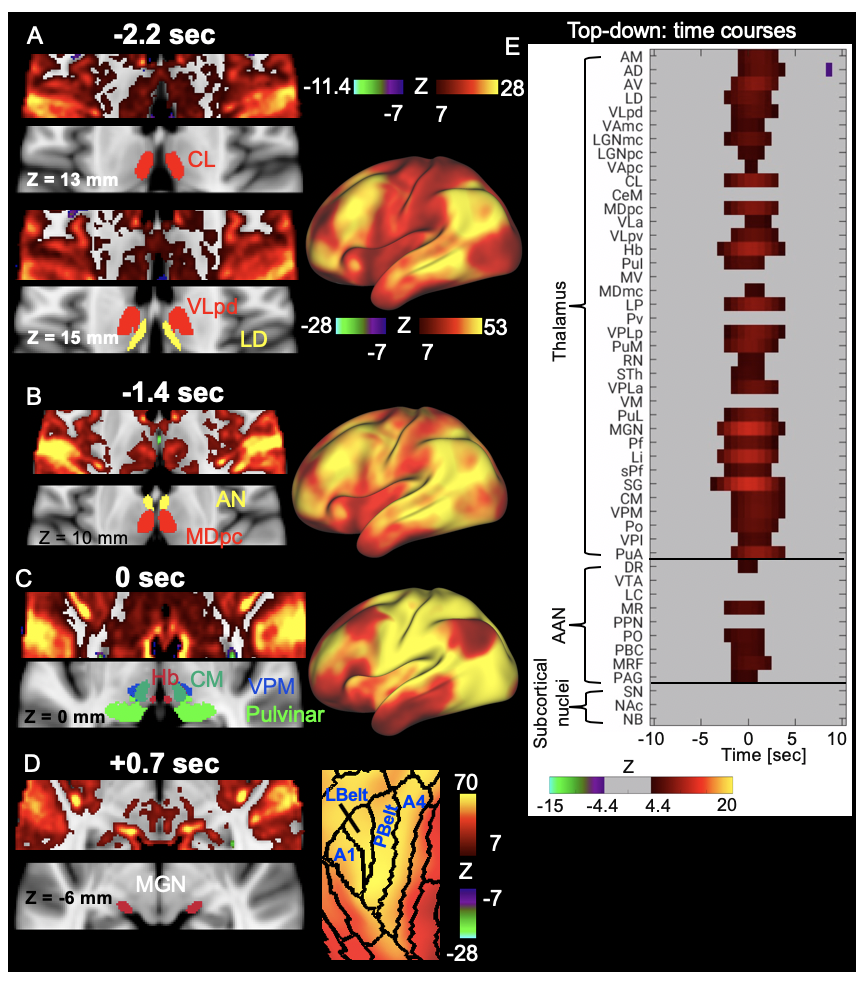

**Fig. S17.**

The relationship between subcortical co-activations and the top-down propagation. The location of the thalamic nuclei was based on the Morel Atlas (6). The time is with respect to the global mean peak. (A-D) The thalamic co-activations at different phases of the top-down propagations show a correspondence with the cortical changes. (A) The co-activation of the DMN is associated with the thalamic co-activations in the CL, VLpd and LD. (B) The co-activation of the DMN is associated with the thalamic co-activations in the AN and MDpc. (C) The co-activation of the sensory/motor is associated with the thalamic co-activations in the Hb, CM, VPM, and pulvinar. (D) The co-activations of MGN is associated with the auditory co-activation (A1, Lbelt, Pbelt, A4). (E) The temporal dynamics of all the subcortical regions. The Z-score time courses were averaged within 37 thalamic regions of interest (ROIs) defined by the Morel’s atlas and 9 brainstem ROIs defined by the ANN atlas, as well as the three ROIs (SN, NAc, NB) we defined by combining our results with brain atlas (7–9). The time courses within each group of ROIs were sorted according to their values at *t* = -5.8 sec. The q-value generated by FDR corresponding to z-score 4.4 and 7 is 10^-5^ and 7.4x10^-13^ respectively. Abbreviation: MDmc, mediodorsal nucleus magnocellular part. MDpc, mediodorsal nucleus parvocellular part. MV, medioventral nucleus. CL, central lateral nucleus. CeM, central median nucleus. CM, centre median nucleus. Pv, paraventricular nucleus. Hb, Habenular nucleus. Pf, parafascicular nucleus. sPf, subparafascicular nucleus. PuM, medial pulvinar. PuI, inferior pulvinar. PuL, lateral pulvinar. PuA, anterior pulvinar. LP, lateral posterior nucleus. MGN, medial geniculate nucleus. SG, suprageniculate nucleus. Li, limitans nucleus. Po, posterior nucleus. LGN, lateral geniculate nucleus. VPLa, ventral posterior lateral nucleus anterior part. VPLp, ventral posterior lateral nucleus posterior part. VPM, ventral posterior medial nucleus. VPI, ventral posterior inferior nucleus. VLa, ventral lateral anterior nucleus. VLpd, ventral lateral posterior nucleus dorsal part. VLpv, ventral lateral posterior nucleus ventral part. VAmc, ventral anterior nucleus magnocellular part. VApc, ventral anterior nucleus parvocellular part. VM, ventral medial nucleus. AD, anterior dorsal nucleus. AM, anterior medial nucleus. AV, anterior ventral nucleus. LD, lateral dorsal nucleus. AN, anterior nucleus. STh, subthalamic nucleus. DR, dorsal raphe. VTA, ventral tegmental area. LC, locus coeruleus. MR, median raphe. PPN, pedunculopontine nucleus. PO, pontis oralis. PBC, parabrachial complex. MRF, midbrain reticular formation. PAG, periaqueductal gray. NAc, nucleus accumbens. NB, nucleus Basalis. SN, substantia Nigra. A4, auditory 4 complex. PBelt, parabelt complex. LBelt, lateral belt complex. A1, primary auditory cortex. SM, sensory/motor. DMN, default mode network.

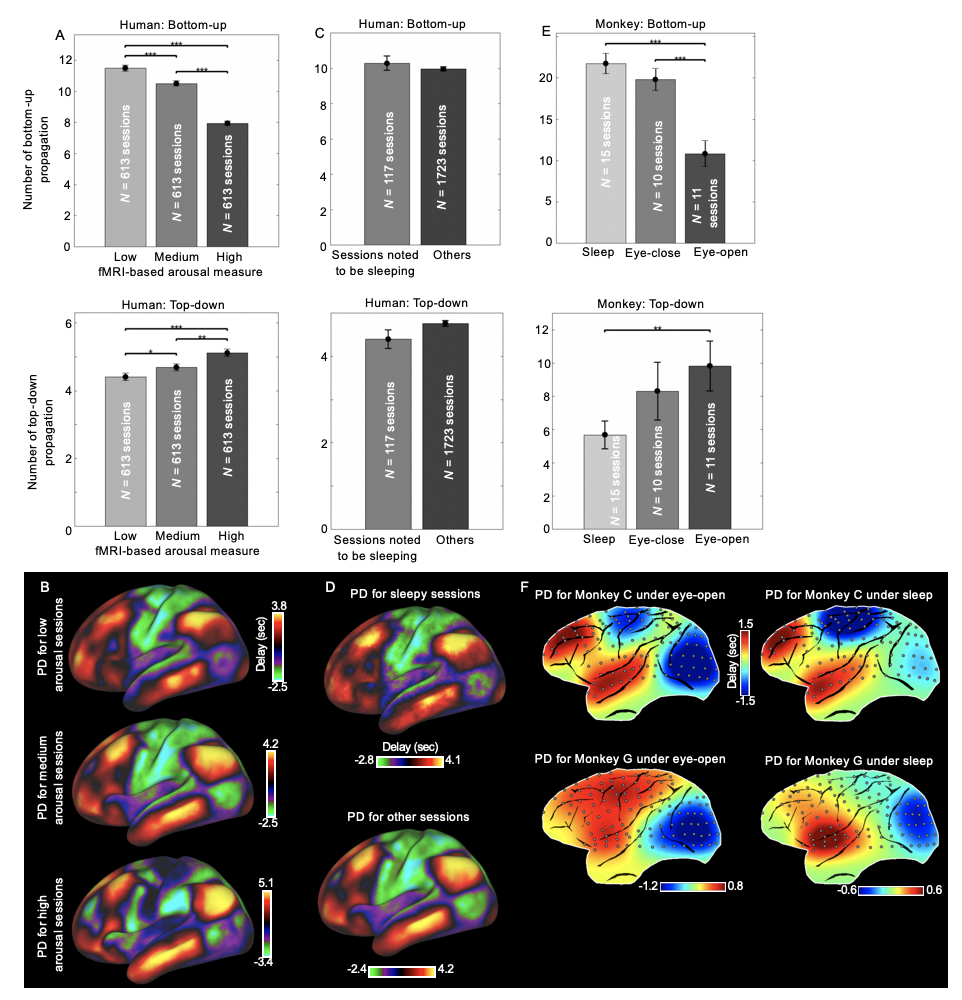

**Fig. S18.**

The low arousal level is related with less top-down and more bottom-up propagation activity. (A) More top-down propagation (bottom panel) and less bottom-up (top panel) propagation is significantly observed in three groups of human rsfMRI sessions with increasing arousal levels as measured by an fMRI-based arousal index (10). The three equal groups of sessions with low, medium, high arousal levels were formed by sorting all the four rsfMRI sessions from 460 subjects based on their arousal level. (B) The principal delay profile is extracted for each group in (A) and all of them showed a strong contrast between the sensory/motor areas and the default mode network, similar with that extracted using all sessions in Fig. 2C. (C) The same trend is also observed in a group of rsfMRI sessions (117 sessions) where subjects were noted to be sleeping during scanning compared to other rsfMRI sessions. (D) The principal delay profile is extracted from the group of sessions noted to be sleeping and the other sessions in (C) respectively and both of them showed a strong contrast between the sensory/motor areas and the default mode network, similar with that extracted using all sessions in Fig. 2C. (E) The number of the top-down propagation and the bottom-up propagation in the ECoG gamma powers shows a similar and significant trend across the eyes-open, eyes-closed, and sleep sessions using ECoG gamma-band powers concatenated from all of the four monkeys. (F) The principal delay profile is extracted under eyes-open or sleep conditions from the ECoG gamma powers for monkey C and monkey G respectively. The ECoG data under both eyes-open and sleep conditions were recorded only in monkey C and monkey G. All of them showed a strong contrast between the sensory/motor areas and the high-order regions, similar with that extracted under eyes-closed conditions in Fig. 3A. Note that the principal delay profile for the human rsfMRI signals in (C-D) is the first principal component; the principal delay profile for the Monkey C under eyes-open condition and sleep condition is the second and the first principal component respectively; the principal delay profile for the Monkey G under eyes-open condition and sleep condition is the first and the second principal component respectively. Each session is 15 minutes and 25 minutes in length for the human rsfMRI data (A-D) and ECoG gamma band powers (E-F) respectively. Error bars represent the standard error of the mean (SEM). Asterisks represent the level of significance: *: 0.01 < *p* $\leq$ 0.05; **: 0.001 < *p* $\leq$ 0.01; ***: *p* $\leq$ 0.001.

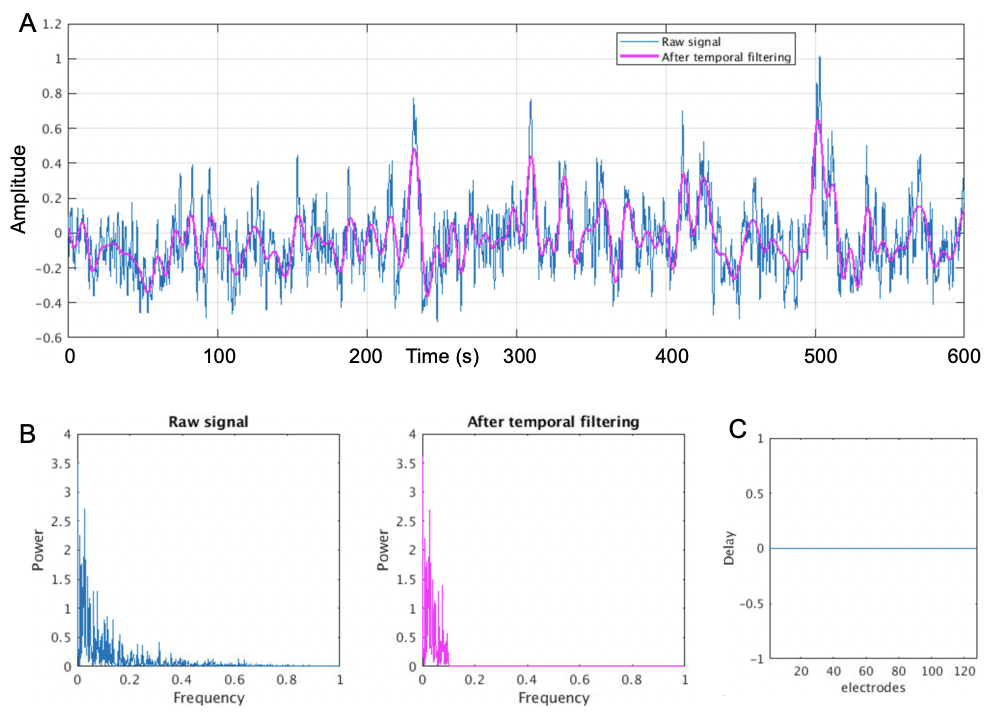

**Fig. S19.**

The performance of the temporal filtering used in the analyses. (A) An example time series of the gamma-band power of monkey ECoG signal (blue) and its temporally filtered (<0.1 Hz) version. (B) The power spectrum of the gamma power signal and the filtered signal. (C) The phase delay between the unfiltered and filtered signals at each electrode.

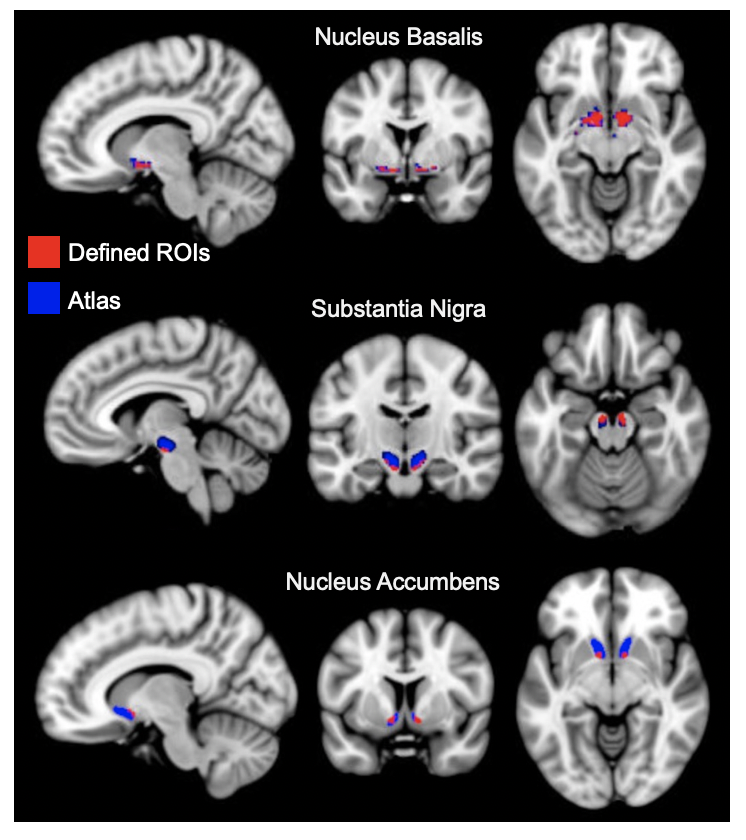

**Fig. S20.**

The three subcortical ROIs defined in our analysis (red) and the corresponding structures defined by atlases (blue).
